# Emerging harmful algal blooms caused by distinct seasonal assemblages of the toxic diatom *Pseudo-nitzschia* in Narragansett Bay, RI, USA

**DOI:** 10.1101/2021.08.18.456122

**Authors:** Alexa R. Sterling, Riley D. Kirk, Matthew J. Bertin, Tatiana A. Rynearson, David G. Borkman, Marissa C. Caponi, Jessica Carney, Katherine A. Hubbard, Meagan A. King, Lucie Maranda, Emily J. McDermith, Nina R. Santos, Jacob P. Strock, Erin M. Tully, Samantha B. Vaverka, Patrick D. Wilson, Bethany D. Jenkins

## Abstract

The diatom *Pseudo-nitzschia* produces the neurotoxin domoic acid (DA) that bioaccumulates in shellfish, causing illness in humans and marine animals upon ingestion. In 2017, high levels of DA in shellfish meat closed shellfish harvest in Narragansett Bay (NBay), Rhode Island for the first time in history, although abundant *Pseudo-nitzschia* have been observed for over 50 years. What caused these events is unknown: whether an environmental factor altered endemic *Pseudo-nitzschia* physiology or new DA-producing strain(s) were introduced. To investigate, we conducted weekly sampling from 2017-2019 to compare with 2016 precautionary closure and 2017 closure samples. Particulate DA was quantified by highly sensitive LC-MS/MS and correlated with environmental metadata. *Pseudo-nitzschia* were identified using high-throughput rDNA sequencing, yielding a detailed understanding of distinct seasonal multi-species assemblages. Low DA was detected throughout 2017-2019, except in recurring peaks in the fall and early summer. Fall DA peaks contained toxigenic species (*P. pungens* var. *pungens, P. multiseries, P. calliantha,* and *P. subpacifica*) as well as a novel *P. americana* taxon. Fewer species were present during summer DA peaks including toxigenic *P. multiseries, P. plurisecta,* and *P. delicatissima.* Most 2017 closure samples contained *P. australis.* Our data showed *P. australis* as infrequent but particularly concerning. Recurring *Pseudo-nitzschia* assemblages were driven by seasonal temperature changes and DA correlated with low dissolved inorganic nitrogen. Thus, the NBay closures were likely caused by resident assemblages dependent on nutrient status as well as the episodic introductions of species that may be a result of oceanographic and climactic shifts.

## Introduction

Phytoplankton photosynthesis sustains oceanic food webs and generates nearly half of photosynthetically fixed carbon on Earth (Falkowski 1994). Despite their key roles in ocean ecosystems, some phytoplankton generate toxins that can be harmful to animal and human health, including diatoms in the genus *Pseudo-nitzschia. Pseudo-nitzschia* can cause harmful algal blooms (HABs) in locations across the globe (Hasle 2002; Bates et al. 2018; Anderson et al. 2019). *Pseudo-nitzschia* spp. produce domoic acid (DA), a water-soluble, neuroexcitatory toxin that serves as an agonist of glutamate receptors in the central nervous system of animals (Bates et al. 1989). DA disrupts marine food webs as it bioaccumulates in vectors such as copepods, anchovies, and shellfish, causing mortality in sea birds, sea lions, and whales (Work et al. 1993; Lefebvre et al. 1999; Fire et al. 2010; Tammilehto et al. 2015). DA ingestion in humans leads to amnesic shellfish poisoning (ASP) with symptoms such as seizures and permanent short-term memory loss. Deaths from ASP resulted from the first known DA event in 1987 on Prince Edward Island, Canada (Bates et al. 1989). However, there may be hidden ASP fatalities and cases due to unconsidered differential diagnosis (Bates et al. 2018). Long-term exposure to low DA concentrations via high shellfish diets may also be harmful, causing diminished memory (Grattan et al. 2018). In addition to human health impacts, DA events have far-reaching effects on coastal communities including public panic and massive income losses in shellfish sales (Hoagland et al. 2002).

Although DA shellfish closures frequently occur along the Gulf of Mexico and the Pacific coasts of the US and Canada, they have not occurred in northeastern US waters until very recently. The first northeastern US closure due to DA exceeding regulatory action limits in shellfish meat was in the Gulf of Maine in September 2016 which resulted in the market recall of shellfish product (Bates et al. 2018). This 2016 closure in Maine was followed by precautionary closures in Massachusetts and Rhode Island (RI) in October 2016 (Fig. S1A) (Bates et al. 2018). Less than a year later in March 2017, a second northeastern closure occurred, which was limited to only Narragansett Bay (NBay), RI with DA levels in shellfish meat exceeding the 20 µg DA/g regulatory limit (Fig. S1B) (Bates et al. 2018; 2019a). Subsequent closures followed in fall 2017 and winter 2018 in eastern and western Maine (Bates et al. 2018). Overall, DA closures due to toxic *Pseudo-nitzschia* spp. are an emergent problem in the northeastern US.

NBay is a unique platform for gaining insights into causative drivers of *Pseudo-nitzschia* HABs. It is the location of one of the longest running plankton time-series in the world, the NBay Long-Term Plankton Time Series (NBPTS, Fig. 1) (https://web.uri.edu/gso/research/plankton/, https://www.nabats.org/). The NBPTS data show *Pseudo-nitzschia* spp. have been present for over 50 years and often at high cell abundances (Fig. 2). Species within the *Pseudo-nitzschia* genus differ in their ability to produce toxin, making species identification important for HAB monitoring (Bates et al. 2018). However, little is known about the *Pseudo-nitzschia* species composition in NBay. *Pseudo-nitzschia* spp. are enumerated by the NBPTS using light microscopy which is insufficient for species delineation as many are cryptic (Lundholm et al. 2003, 2006; Orsini et al. 2004; Amato et al. 2007; Quijano-Scheggia et al. 2009). Therefore, it is difficult to assess the presence or absence of toxigenic species in historical NBay data. Electron microscopy has been used on selected samples from NBay and nearby waters and has identified six toxic species that can co-occur: *P. delicatissima, P. fraudulenta, P. multiseries, P. pseudodelicatissima, P. pungens* var. *pungens,* and the typically offshore *P. seriata* (Hargraves and Maranda 2002). Given the recent DA closures despite the persistence of *Pseudo-nitzschia* spp. and DA monitoring since the early 1990s, it is critical to understand the ecological species dynamics and DA production of *Pseudo-nitzschia* spp. in NBay.

**Fig 1.**
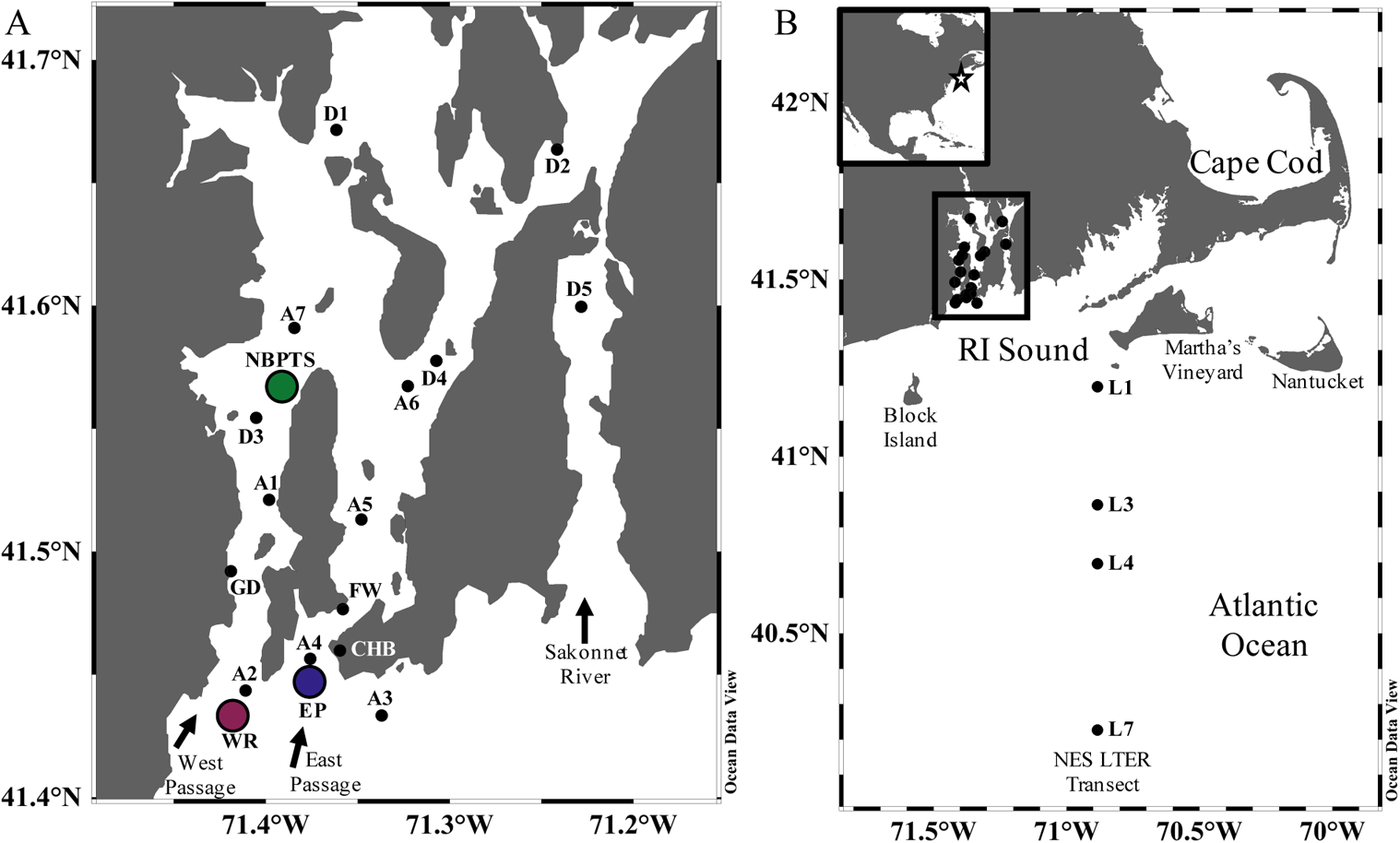
Sampling sites within Narragansett Bay (NBay), RI, USA and offshore Northeast U.S. Shelf (NES) Long-Term Ecological Research (LTER) cruise stations. **[A]** Sample locations (n = 18) within NBay including the Long-Term Plankton Time Series (NBPTS) at the green point, University of RI Graduate School of Oceanography dock (GD), Whale Rock (WR) at the purple point, East Passage (EP) as the blue point, Castle Hill Beach (CHB), and Fort Wetherill (FW), and sites sampled by RI harmful algal bloom (HAB) monitoring in March 2017 (D1 - D5) and sites sampled with vertical net tows (A1 - A7). **[B]** Inset shows overview of sampling site locations in North America. The location of the within-NBay sites are outlined by the black box. Select stations were sampled along the routine NES LTER transect (L1, L3, L4, L7). Figure made in Ocean Data View (65).

**Fig 2.**
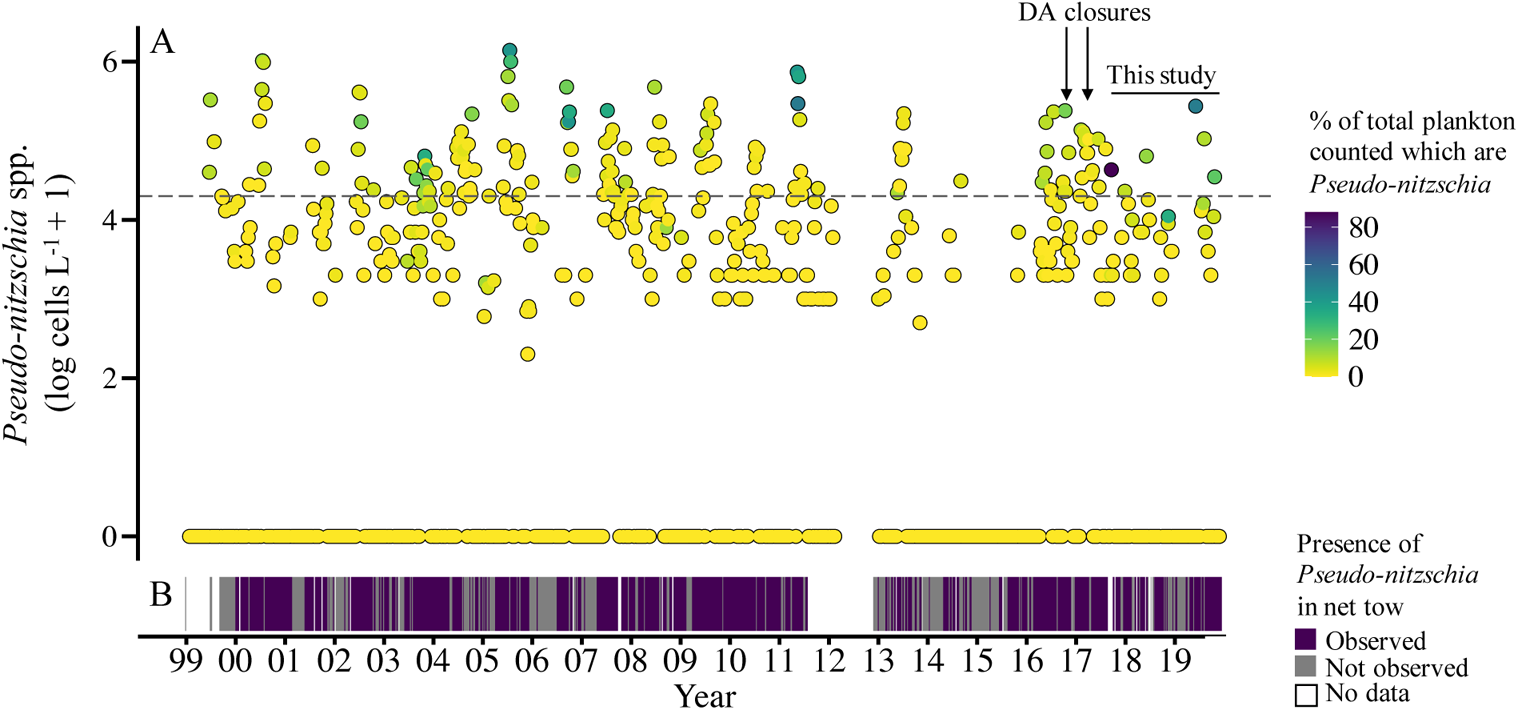
Twenty-year record of the presence of *Pseudo-nitzschia* spp. in Long-Term Plankton Time Series (NBPTS) samples in Narragansett Bay, RI **[A]** Counts of live *Pseudo-nitzschia* spp. cells observed at the NBPTS site from approximately weekly sampling from Jan 1999 - Nov 2019 (n = 1003). Values are Log_10_-transformed, with +1 cell to all values to create non-zero values in the transformation. Surface samples were used in 2000, 2001, and 2008 - 2019, and mixed surface and depth (∼ 7 m) samples were used in 1999 and 2002 - 2007. No sampling occurred from March 2012 – December 2012. The horizontal dotted line is the log-transformed RI HAB action threshold of 20,000 cells L^-1^. The line annotated “This study” is the timeframe that samples were taken for DNA sequencing and particulate domoic acid (pDA).

Arrows indicate the 2016 precautionary closure and 2017 closure due to DA, respectively. The color of the points indicates the percentage of total plankton community counts that *Pseudo-nitzschia* spp. represent. **[B]** Data from 20 µm vertical net tow samples (∼ 7 m bottom depth) corresponding to count data presented in A; each line represents whether *Pseudo-nitzschia* spp. were observed or not (n = 920). Net tow data unavailable from September 2011 – March 2012.

Several hypotheses can explain the recent DA closures in NBay. It is possible that a particularly toxigenic species of *Pseudo-nitzschia* recently became present or much more prevalent. Alternatively, or in addition, environmental dynamics may have shifted to increase toxin production by resident species. Toxigenic species regulate DA production in response to a variety of factors including silicate limitation (Bates et al. 1996), phosphate limitation (Pan et al. 1996), different nitrogen sources (Auro and Cochlan 2013), and in multifactor combinations such as silicate limitation and pCO_2_ (Tatters et al. 2012). Thus, one or more environmental factors may have stimulated DA production in NBay.

To delineate the relationships between the environment, *Pseudo-nitzschia* species composition, and DA production, we conducted weekly time series sampling at several sites in NBay from 2017-2019 (Fig. 1). Offshore sampling transects were used to examine linkages between NBay species assemblages and those on the Northeast US Shelf (NES). Particulate DA (pDA) from plankton samples (> 5 µm size) was directly quantified with tandem mass spectrometry coupled with liquid chromatography (LC-MS/MS). Typically, in HAB monitoring, LC-MS/MS is not employed in high frequency sampling but instead used as confirmation for indirect measurements, such as ELISA-based methods or the Scotia Rapid Test. These frequent and highly sensitive pDA measurements provided an unprecedented understanding of toxin dynamics, especially of levels below closure concerns.

*Pseudo-nitzschia* species composition was delineated using new molecular barcoding methodology targeting the 18S – 5.8S rDNA internal transcribed spacer region 1 (ITS1). This high-throughput sequencing approach allowed for species to be catalogued at high temporal resolution, and circumvented labor intensive and time-consuming electron microscopy. In addition, we improved upon previously used molecular methodologies that relied on ITS1 amplification with fragment size polymorphism to distinguish a select number of *Pseudo-nitzschia* species (Hubbard et al. 2008, 2014). We targeted the ITS1 region with a custom primer designed from an updated database of global *Pseudo-nitzschia* rDNA sequences to yield unique sequences from at least 41 different species. Overall, comparing pDA production, species composition in toxic, non-toxic, and closure samples along with environmental variables informed our ecological understanding of *Pseudo-nitzschia* HABs in NBay. We can begin to predict their temporal patterns and magnitude, with potential implications for the larger region of the Atlantic Northeast. Ultimately, our results will inform forecasting HAB models and management decisions of RI shellfish harvest.

## Materials and Methods

### Sample sites

NBay is a temperate estuary in Rhode Island, USA with seasonally reoccurring phytoplankton blooms in the winter with smaller blooms in the summer (Oviatt 2004). The Providence River contributes inflow of freshwater and urban nutrients in the north, and Atlantic Ocean water is introduced from RI Sound in the south into the East and West passage mouths of NBay. From September 2017 – November 2019 (referred to as the “study period” in figures and text), surface seawater samples were collected weekly or bi-weekly at multiple sites in NBay (Fig. 1, Table S1, Schlitzer 2002). Weekly sampling was conducted at the mid-Bay Narragansett Bay Long-Term Plankton Time Series (NBPTS) site (41.57 N, −71.39 W). The Whale Rock (WR) site at the mouth of West Passage (41.34 N, −71.42 W) is the location of the URI GSO NBay Fish Trawl Survey and samples were collected at this location starting in October 2018.

**Table 1.**
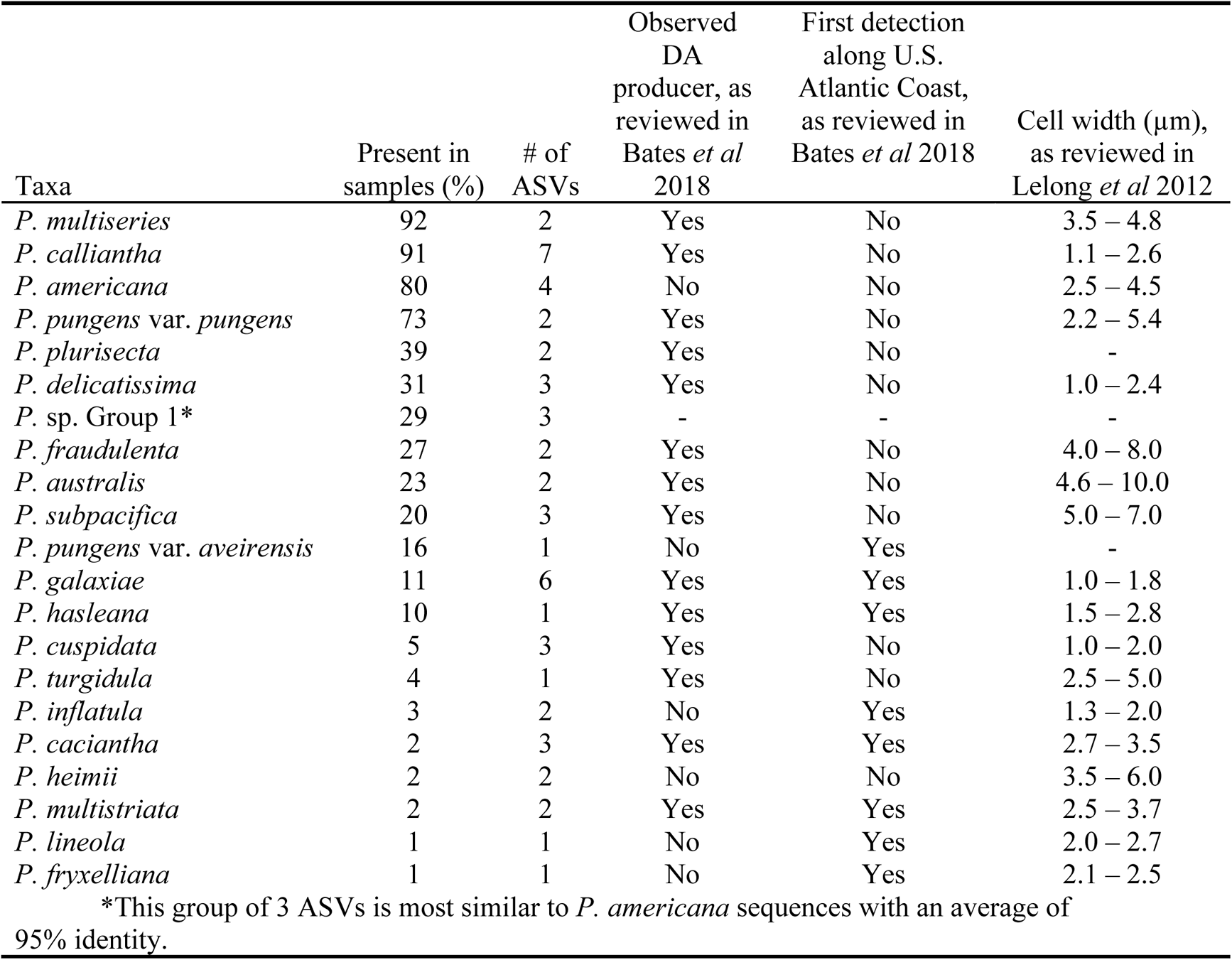
Summary of the 53 *Pseudo-nitzschia* spp. amplicon sequence variants (ASVs) recovered from NBay 2016 - 2019 samples, including offshore Northeast Shelf Long-Term Ecological Research (NES LTER) cruise samples (n = 192), that are present at 1% or higher relative abundance in individual samples out of all *Pseudo-nitzschia* ASVs.

The EP site (41.45 N, −71.38 W) is at the East Passage mouth and a location for RI HAB monitoring. During spring and fall time periods when *Pseudo-nitzschia* spp. cells were observed or pDA was detected, additional sampling on the same day was conducted at the NBPTS and WR sites along with EP. Otherwise, the frequency of each sample is about once a week.

Additional shore-based sites also supplemented sampling: Castle Hill Beach (EP shore access), Fort Wetherill (EP shore access), and GSO Dock (NBPTS alternative site) (Table S1). Vertical net tow samples from various sites in NBay were also collected by L. Maranda in fall 2017, summer - fall 2018, and spring - fall 2019, with corresponding pDA Scotia Rapid Tests and *Pseudo-nitzschia* spp. cell counts, from an average of 1.7 m^3^ water concentrated and zooplankton removed by filtering across a 150 µm mesh. These samples are referred to as “net tow samples” in text. Surface seawater samples from the 2017 closure were collected by RI DEM on March 13, 2017 from five sites in mid-Bay and sent to K. Hubbard for DNA extraction; however, all these sites were north of the March 2017 shellfish harvest closure area. In addition, four NES LTER transect cruises onboard the R/V *Endeavor* provided offshore surface samples in 2018 and 2019 (Table S1).

### Environmental metadata

Surface seawater temperature and salinity were measured using YSI Sondes (YSI Inc. / Xylem Inc., Yellow Springs, OH, USA): EXO Sonde from May - August 2018 at sites outside the NBPTS and WR, ProDSS starting September 2018, and 6920 V2 for weekly NBPTS and Fish Trawl Survey samples from October 2018 - September 2019. Data from the NBPTS are available: https://web.uri.edu/gso/research/plankton/data/, as are 2017 Fish Trawl Survey data: https://web.uri.edu/fishtrawl/data/. Fish Trawl Survey data from 2018 – 2019 at WR were acquired from the fish trawl personnel. Extracted chlorophyll *a* measurements were determined by vacuum filtering triplicate surface seawater onto GF/F filters (0.6 – 0.8 µm particle retention; Whatman n.k.a. Cytiva, Marlborough, MA, USA). Filters were extracted in 90% acetone for 24 hours in the dark at −20 °C. After 24 hours, samples were equilibrated at room temperature for 20 minutes, vortexed, and decanted for fluorometric reading in a 10AU fluorometer (Turner Designs, Inc., San Jose, CA, USA) for May - July 2018 and NBPTS samples or a Trilogy fluorometer (Turner Designs, Inc., San Jose, CA, USA) for samples August 2018 - October 2019. Chlorophyll *a* was determined as in the Turner Designs manual by subtracting values from samples acidified with three drops of 10% hydrochloric acid (2019b).

Samples for dissolved nutrient analysis were filtered through 0.2 μm polyethersulfone filters (Sterlitech, Kent, WA, USA) and frozen at −20 °C until analysis on a Lachat QuickChem 8500 (Hach, Loveland, CO, USA) at the URI Marine Science Research Facility (Narragansett, RI, USA) or on AA3 (SEAL Analytical, Inc., Mequon, WI, USA) at the University of Washington Nutrient Analysis Facility (Seattle, WA, USA) including duplicate samples for comparison between facilities. The UW AA3 measures nitrate directly, whereas the Lachat nitrate values were calculated from nitrite + nitrate minus nitrite. Any negative Lachat values for ammonium were changed to zero, but all other values were used in analysis. For any analysis showing DIN, the summation of ammonium, nitrate, and nitrite was used.

*Pseudo-nitzschia* spp. cells were enumerated using the methods of the URI GSO NBPTS. Details are provided in the Supplementary Information. *Pseudo-nitzschia* were recorded as observed or not observed in net tow samples (>20 µm). Current and historical NBPTS cell abundance data are available online: https://web.uri.edu/gso/research/plankton/. The action threshold of *Pseudo-nitzschia* spp. cell counts of 20,000 cells L^-1^ is highlighted in some figures, which is from the RI HAB Plan (2020).

### Pseudo-nitzschia spp. ITS1 DNA analysis

Plankton biomass for *Pseudo-nitzschia* spp. DNA identification was collected with a peristaltic pump, passing an average of 270 mL of surface seawater across a 25 mm 5.0 μm polyester membrane filter (Sterlitech, Kent, WA, USA). Widths of some *Pseudo-nitzschia* spp. are < 5.0 μm (Lelong et al. 2012), but the 5µm size pore likely captured chains and horizontally orientated cells and was consistent with pore size used to examine toxicity. Filters were flash frozen in liquid nitrogen and stored at −80 °C until DNA extraction. See Supplemental Information for the DNA extraction protocol and a description of sequencing controls.

The 18S V4 rDNA region was initially amplified using primers specific to diatoms (Zimmermann et al. 2011) however, this highly conserved region is not able to fully delineate *Pseudo-nitzschia* species, but the ITS1 region discriminates between *Pseudo-nitzschia* spp. (Hubbard et al. 2008). To target the *Pseudo-nitzschia* ITS1 we used an existing eukaryotic ITS1 forward primer: 5’ TCCGTAGGTGAACCTGCGG 3’ (White et al. 1990) and designed a custom reverse primer (Pn-ITS1R) against a conserved 5.8S region using 132 *Pseudo-nitzschia* ITS1 sequences (Table S2) (NCBI nucleotide database as of 4/3/2019): 5’ CATCCACCGCTGAAAGTTGTAA 3’. We added MiSeq adapters to the 5’ ends of these primers and the full primers used were: forward 5’ TCGTCGGCAGCGTCAGATGTGTATAAGAGACAGTCCGTAGGTGAACCTGCGG 3’ and reverse 5’ GTCTCGTGGGCTCGGAGATGTGTATAAGAGACAGCATCCACCGCTGAAAGTTGTAA 3’.

*Pseudo-nitzschia* species expected to amplify with these primers are summarized (Table S2). The expected ranges for PCR products were from 235 – 370 bp as the size of the ITS1 region differs for some *Pseudo-nitzschia* taxa. PCR amplification details are provided in the Supplementary Information. Sequence library preparation and 2×250 bp Illumina MiSeq (Illumina, Inc., San Diego, CA, USA) sequencing were performed by the RI Genomics and Sequencing Center (Kington, RI, USA). There were 193 environmental samples sequenced, along with the negative and positive controls (see Supplemental Information), for a total of 196 samples using two sets of MiSeq indices on the same sequencing plate. The raw sequencing reads are available on NCBI’s Short Read Archive (Bioproject PRJNA690940).

Sequences were analyzed with a custom bioinformatics pipeline. llumina MiSeq adapters and primers were trimmed from both read ends using CutAdapt v1.15 (Martin 2011) and input to DADA2 v1.16 (30) to determine amplicon sequence variants (ASVs). Reads lacking ITS1 primer sequences were discarded. Unique ASVs, some differing by one bp, were included in subsequent analysis. ASVs were identified as *Pseudo-nitzschia* taxa using a curated database from NCBI sequences (Table S2), which was also used to design the reverse primer, and the scikit-learn naïve Bayes machine learning classifier (Pedregosa et al. 2011) at default settings in QIIME2 v2020.2 (Bolyen et al. 2019). The scikit-learn naïve Bayes machine learning classifier identified 94 ASVs from environmental samples as *Pseudo-nitzschia* at the species level. Additional *Pseudo-nitzschia* ASVs were assigned using a BLAST pipeline; details are provided in Supplementary Information. All ASVs identified in this study as *Pseudo-nitzschia* are deposited into NCBI GenBank under accession numbers MW447658-MW447770. Read counts were transformed into relative abundance out of total reads assigned to *Pseudo-nitzschia*. If an ASV accounted for < 1% relative abundance in a sample, then it was considered “not present” to avoid potentially spurious results. This removed 60 ASVs from consideration. The remaining 53 ASVs were analyzed in a presence/absence matrix to avoid potential problems from inflating read numbers with cell counts and rDNA copy number variation (Jenkins and Bertin 2021a).

### Domoic acid analysis

At each sampling station, biomass was collected by filtering approximately 2 L of surface seawater across a 47 mm 5.0 μm polyester membrane filter (Sterlitech, Kent, WA, USA). Filters were flash frozen in liquid nitrogen and stored at −80 °C until extraction. Filters were extracted for four hours in 0.1 M acetic acid, vigorously vortexing each hour. Extracts were passed through a 0.2 µm syringe filter directly into a 1.5 mL LC-MS vial for LC-MS/MS analysis on a Prominence UFLC system (Shimadzu, Kyoto, Japan) coupled to a SCIEX 4500 QTRAP mass spectrometer (AB Sciex, Framingham, MA, USA). Mass Spectrometry methodological details are provided in the Supplemental Information. Plankton-associated DA was quantified to ng pDA L^-1^ of filtered seawater using an external calibration curve performed from pure DA standards of increasing concentrations (Sigma-Aldrich, St. Louis, MO, USA), included in each analysis. DA data and environmental metadata are publicly available through the Biology and Chemical Oceanography Data Management Office, BCO-DMO (Jenkins and Bertin 2021b).

DA was also analyzed from in the tissue of six live mussels (RI DEM Scientific Collector’s Permit #435) collected at the GSO dock on June 5, 2019 during a period of increased *Pseudo-nitzschia* cell abundance and elevated pDA (see the Supplementary Information for methodological details). Scotia Rapid Tests were performed for from the offshore A3 site net tow samples on approximately 950 mL of the net tow concentrate.

### Statistical analysis and Visualization

For the principle components analyses (PCA), all samples from any site with pDA measurements were used and included some which were not sequenced. Data were transformed on a log scale using the addition of a constant = 1, except for surface water temperature where the constant 10 was used and for salinity where no constant was needed. Each variable was standardized by subtracting the mean from its value and then dividing by the standard deviation (in factoextra, scale = TRUE) (Kassambara and Mundt 2020). Levene’s test was performed using the car package v3.0.10 in R with command leveneTest() (Fox and Weisberg 2019), one-way ANOVA was run in base R with aov(), and Tukey HSD was performed with TukeyHD() in base R. The function BIOENV (Best Subset of Environmental Variables) through the vegan bioenv() command was used to determine the best model formula from the scaled environmental variables used in the PCA for 127 sequenced NBay samples of the Bray-Curtis distance of the *Pseudo-nitzschia* ASVs (Clarke and Ainsworth 1993).

Similarities between the presence/absence of Jaccard distance of *Pseudo-nitzschia* ASVs across samples were explored with ANOSIM and NMDS. A test for dispersion of groups was performed prior to ANOSIM, to assess reliability of ANOSIM results. Dispersion of groups was tested using command betadisper() in the R package vegan. Then within the same package, permutest() performed on those results with 999 permutations and alpha value of 0.05 was used to assess significance. Only groups lacking a significant group dispersion were carried forward into ANOSIM. Groups investigated included season, sample group, month, year, sample site, closure time, pDA grouping, and if *Pseudo-nitzschia* spp. cells were over or under the action threshold. For seasonal groupings, the northern hemisphere meteorological groupings were used which starts a season on the first of the month in which the respective equinox or solstice begins. Thus, seasons were designated as summer (June, July, and August), fall (September, October, and November), winter (December, January, and February), and spring (March, April, and May). Since many groups had uneven numbers, ANOSIM was preferred to PERMANOVA tests.

ANOSIM was run using anosim() in R package vegan, with 999 permutations and an alpha value of 0.05. NMDS plots of the Jaccard distance (Jaccard 1912) of the presence or absence of the *Pseudo-nitzschia* ASVs which passed through the >1% relative abundance per sample threshold were made to examine similarities and dissimilarities in species composition of samples across sampling sites, seasons, and pDA concentrations. NMDS solutions were not reached by defaults, so number of iterations were increased (trymax = 100) and number of dimensions were raised to 3 (k = 3) with 3-D versions of the plots in Fig. S9. The final stress of the NMDS was 0.0924.

Date was visualized in R v4.0.2 (R Core 2020) in R Studio v1.1.456 (RStudio 2016) with packages: vegan v2.5.6 (Dixon 2003) for group dispersion tests and ANOSIM; ggplot2 v3.3.2 (Wickham 2016) for plots such as line graphs and heatmap; phyloseq v1.32.0 (McMurdie and Holmes 2013) for ASV data set manipulation like subsetting and transformations, and NMDS analysis; factoextra v1.0.7 (Kassambara and Mundt 2020) for principal components analysis (PCA) of data related to pDA; and viridis v0.5.1 for continuous scale coloring (Garnier 2018). Colors were chosen to maximize color blind accessibility (Tol 2018).

## Results

In response to the 2016 precautionary closure and 2017 DA closure, we initiated a time series study of pDA and *Pseudo-nitzschia* spp. composition from September 2017 – November 2019 resulting in over 230 samples collected at various sites within NBay and offshore RI Sound and the Atlantic Ocean (Fig. 1). The majority of samples were collected at the NBPTS site in mid-NBay, Whale Rock (WR) at the West Passage mouth, and East Passage (EP) at its corresponding mouth of NBay (Fig. 1A). At seven additional sites ranging from mid-NBay to offshore, vertical plankton net tows (20 µm) were conducted to examine and compare depth-integrated *Pseudo-nitzschia* spp. assemblages (Fig. 1A). To understand how offshore dynamics may be impacting *Pseudo-nitzschia* spp. and pDA dynamics in NBay, we sampled four sites from four NES Long-Term Ecological Research (LTER) research cruises (Fig. 1B). *Pseudo-nitzschia* spp. composition was also assessed in closure samples: five samples each from the 2016 precautionary closure and the 2017 closure at the NBPTS site and five samples collected by the RI HAB monitoring program in March 2017 at sites in mid to upper NBay (Fig. 1A). No LC-MS/MS pDA measurements were made for these closure samples, as samples were not available. For most samples, light microscopy counts of *Pseudo-nitzschia* spp. cells at the genus level were recorded, extracted chlorophyll *a* was used to evaluate total phytoplankton biomass, and filtered seawater was analyzed for inorganic nutrient concentrations.

### Historical and contemporary Pseudo-nitzschia spp. abundance in NBay

From 1999 to 2019, *Pseudo-nitzschia* spp. abundance at the NBPTS site ranged from unobserved to 1,389,000 cells L^-1^ (Fig. 2A). Cell abundances exceeding the RI HAB monitoring program action threshold of 20,000 cells L^-1^ (2020) occurred annually except in 2015 and comprised approximately 16% of counted samples. Elevated *Pseudo-nitzschia* spp. abundance was observed in summer and fall, and *Pseudo-nitzschia* cells made up over 50% of the total enumerated plankton community in May 2011, September 2017, and June 2019 (Fig. 2A). In addition to these weekly direct plankton counts in 1 mL of seawater, the presence and absence of species were recorded from higher volume net tows. While *Pseudo-nitzschia* spp. typically represented a low fraction of the total plankton cells enumerated in 1 mL surface seawater (averaging 1%) and were detected in less than half of 1 mL count samples, *Pseudo-nitzschia* spp. were consistently detected in over 68% of the high volume NBPTS net tow samples (Fig. 2B), thus indicating *Pseudo-nitzschia* spp. were frequently present in NBay, but at concentrations less than 1,000 cells L^-1^.

*Pseudo-nitzschia* spp. abundance was uncoupled from peaks in phytoplankton biomass as measured by chlorophyll *a* concentration, demonstrating that *Pseudo-nitzschia* spp. bloom at distinct times from other phytoplankton (Fig. 3A, Fig. 3B). During our time series study period from September 2017 – November 2019, *Pseudo-nitzschia* spp. abundance and seasonal patterns were similar across the NBPTS site and the East and West entrances of NBay (Fig. 3B). Peak *Pseudo-nitzschia* spp. cell abundance occurred in the summers, and abundance was elevated in two of the three years of fall samples (Fig. 3B). Cell counts exceeded the action threshold annually, with the exception of the EP site in 2017 when only three samples were collected (Fig. 3B). Abundance ranged from unobserved to 148,000 cells L^-1^ at WR and to 125,000 cells L^-1^ at EP (Fig. 3B).

**Fig 3.**
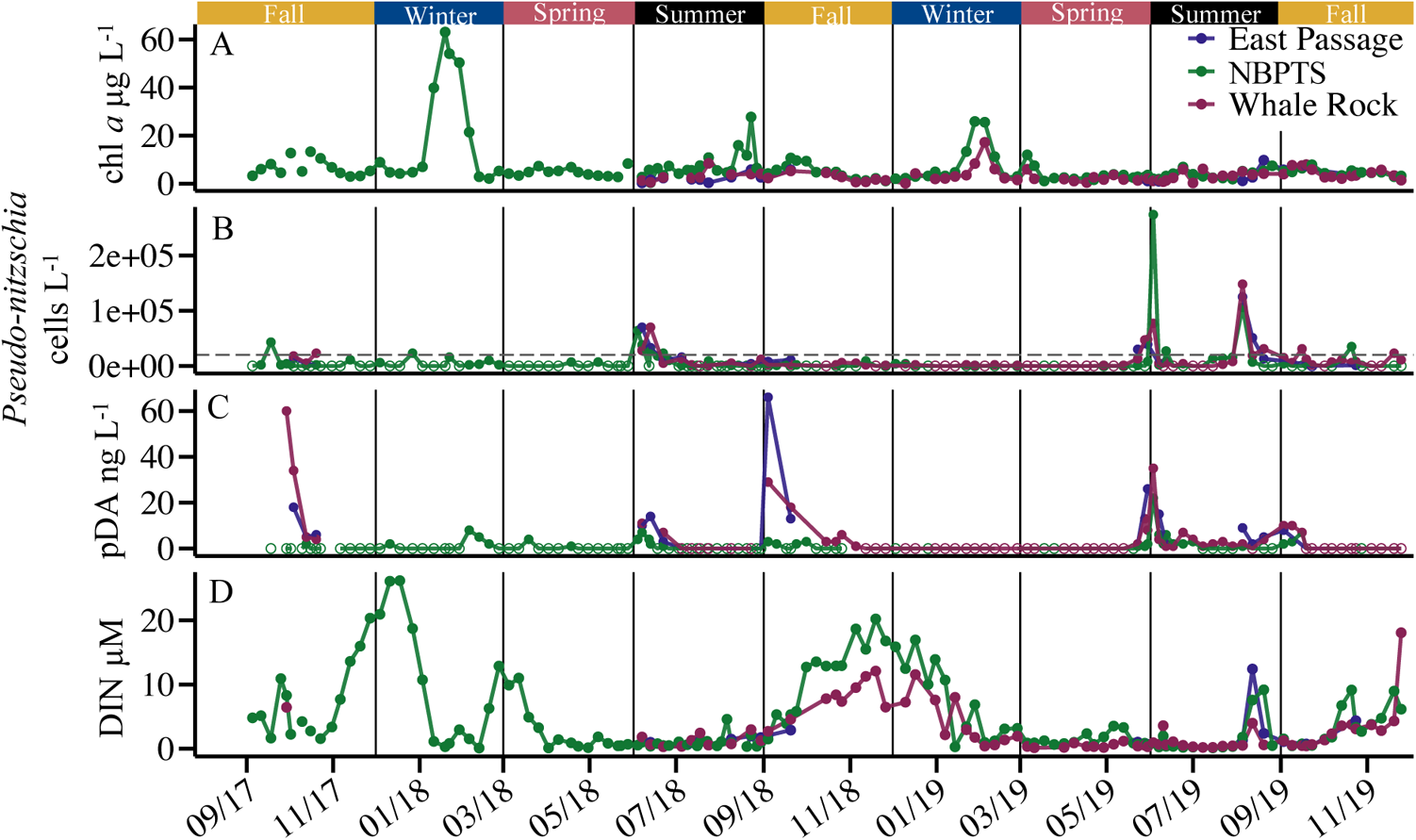
Comparison of total chlorophyll *a* concentration, *Pseudo-nitzschia* spp. cell counts, particulate domoic acid (pDA), and dissolved inorganic nitrogen (DIN) at the most frequently sampled sites during the study period. The East Passage (EP) site was usually sampled in the summer and fall only. **[A]** Trends of chlorophyll *a* concentration across time at the EP site (n = 22), the Long-Term Plankton Time Series (NBPTS) site (n = 131), and Whale Rock (WR) site (n = 69). **[B]** *Pseudo-nitzschia* spp. cell counts across time at EP (n = 26), NBPTS (n = 141), and WR (n = 73). The dotted line is the 20,000 cells L^-1^ action threshold for RI HAB monitoring. Open circles denote the absence of cells. **[C]** Patterns of pDA measured across time at EP (n = 26), NBPTS (n = 131), and WR (n = 72). Open circles denote no pDA was detected. **[D]** DIN, which is the summation of nitrite, nitrate, and ammonium measured, across time at EP (n = 23), NBPTS (n = 136), and WR (n = 71).

### Seasonal patterns of pDA

Our time series data revealed extended periods of low or undetectable pDA concentrations in NBay with distinct seasonal peaks in September, October, May, and June (Fig. 3C, Fig S2). During the late spring and early summer of 2018 and 2019, pDA concentrations were similar at all three sites and corresponded to elevated *Pseudo-nitzschia* spp. cell counts (Fig. 3C, Fig. 3B). In contrast, in the fall, elevated pDA occurred with low *Pseudo-nitzschia* spp. abundance (Fig. 3C, Fig. 3B). Additionally, in the fall of 2017 and 2018, pDA peaks occurred at both the East and West NBay entrances, but not at the mid-Bay NBPTS site (Fig 3C). The NBay entrances also had the highest pDA concentrations measured during our time series with 60 ng pDA L^-1^ at WR in October 2017 and 66 ng pDA L^-1^ at EP in September 2018 when *Pseudo-nitzschia* spp. cell counts were below the action threshold (Fig. 3B, 3C). Despite uncoupling between pDA and *Pseudo-nitzschia* spp. cell counts observed in the fall, there was an overall linear relationship between *Pseudo-nitzschia* spp. cell abundance and pDA concentration at any site sampled during the study period; however, great variability limits the predictive power of the linear model (R^2^ = 0.19, p = 8.08 x 10^-16^, Fig. S3). Although the pDA measured during our study period did not occur at levels triggering concomitant shellfish harvest closures, DA was detected in the tissues of all six mussels sampled on June 5, 2019, during a period of increased *Pseudo-nitzschia* spp. cell numbers and elevated pDA concentrations. Toxin ranged from 0.4 - 4 ng DA/g shellfish meat, and the highest concentration of 4 ng DA/g was well below the mandatory closure level of 20 µg DA/g shellfish meat (2019a).

### Toxin and low dissolved nitrogen concentrations

A principal component analysis (PCA) was used to correlate the chemical and biological properties of all samples collected during this study (Fig. 4). Data were log-transformed and standardized and included pDA concentrations, *Pseudo-nitzschia* spp. cell counts, chlorophyll *a* concentrations, dissolved nutrient concentrations [inorganic nitrogen (DIN): ammonium, nitrite, nitrate), inorganic phosphorus (DIP), and inorganic silicate (DSi)], surface seawater salinity and temperature. Nitrate, ammonium, and nitrite contributed most to the first component, while temperature, DIN:DSi, and DIN:DIP primarily contributed to the second component (Fig. 4A). Temperature, as expected, positively correlated with summer and fall samples and negatively correlated with winter and spring samples (Fig. 4B). *Pseudo-nitzschia* spp. cell counts and pDA concentrations contributed little to variability in the whole dataset, and were correlated with each other suggesting an overarching relationship that is supported by the linear model (Fig. 4, Fig. S3). *Pseudo-nitzschia* spp. cell counts and pDA concentrations showed an inverse relationship to chlorophyll *a* concentrations, nitrate, and ammonium (Fig. 4C). Nitrate and ammonium were negatively correlated with a large cluster of spring, summer, and fall samples in which pDA was detected (Fig. 4C). Notably, increased pDA concentrations were synchronous with low DIN concentrations throughout the time series (Fig. 3C, Fig. 3D) and 82% of samples with detectable pDA occurred when DIN was less than 5 µM (Fig. S4).

**Fig 4.**
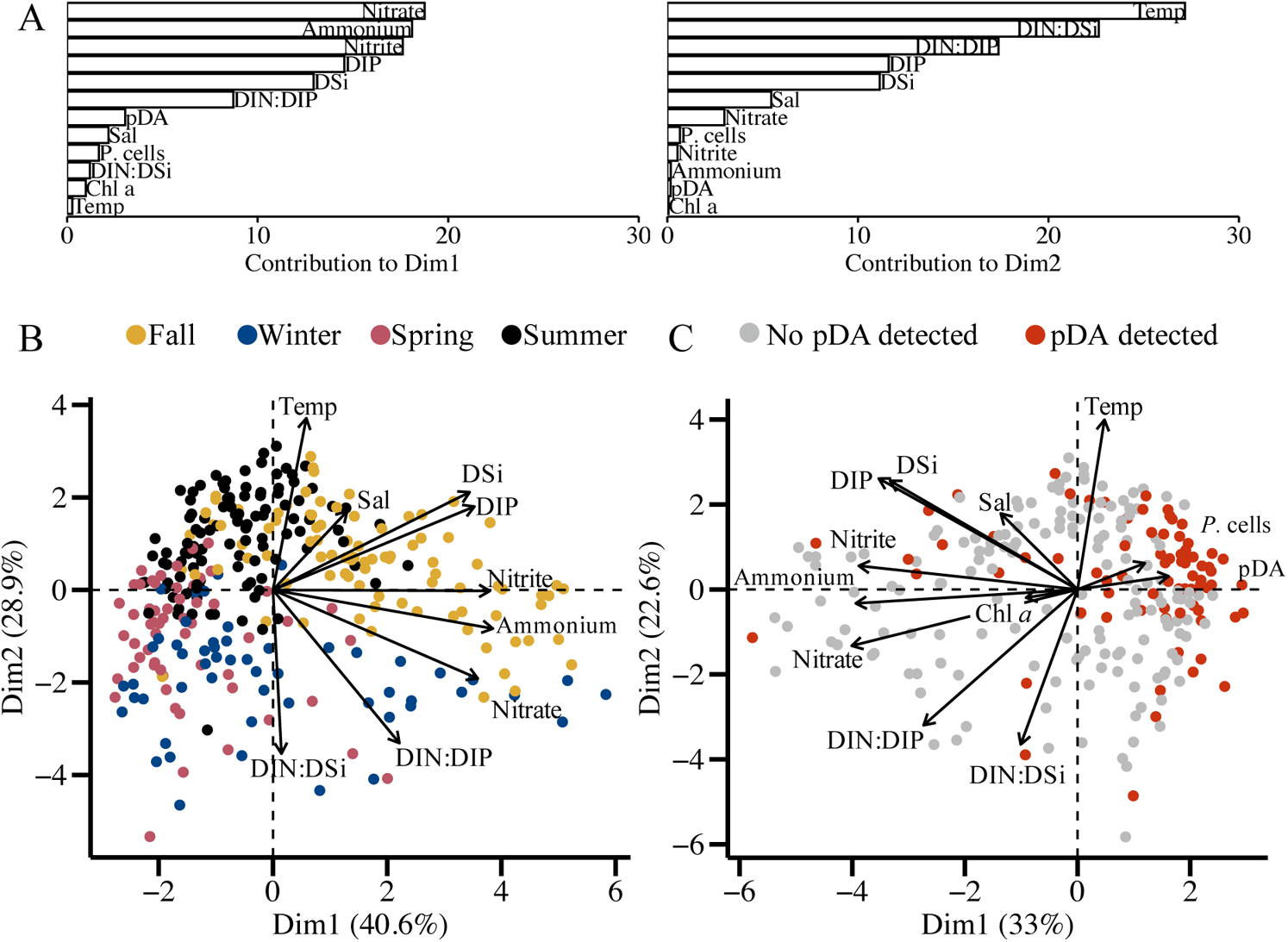
Principal component analysis (PCA) biplot of all surface seawater samples from Narragansett Bay, RI during the study period from any sampling site, excluding NES LTER stations, with full portfolio of physical and chemical parameters log transformed and standardized. Parameters include surface seawater salinity (Sal), surface seawater temperature (Temp), chlorophyll *a* concentration (Chl *a*), dissolved inorganic silicate (DSi), dissolved inorganic phosphorus (DIP), nitrate, nitrite, ammonium, and nutrient ratios related to the summation of those nitrogen sources as dissolved inorganic nitrogen (DIN) of DIN:Si and DIN:P, *Pseudo-nitzschia* spp. cell abundance (*P.* cells), and particulate domoic acid (pDA) concentration. **[A]** Bar chart of the contribution of each variable to Dimension 1 and Dimension 2 corresponding to the PCA shown in C. **[B]** Samples colored by the season sampled, from September 2017 – November 2019, excluding Chl *a, P.* cells, and pDA (n = 312). **[C]** Samples colored by whether pDA was detected (red) or not detected (grey) by LC-MS/MS, and including Chl *a, P.* cells, and pDA (n = 238).

### Pseudo-nitzschia spp. delineated with ITS1 rDNA amplicon sequence variants

The ITS region is one of the most distinctive genetic identifiers for *Pseudo-nitzschia* spp., revealing intraspecific diversity which has implications for toxin production (Lundholm et al. 2003; Orsini et al. 2004; Amato et al. 2007; Casteleyn et al. 2008; Hubbard et al. 2008; Kaczmarska et al. 2008). We used high-throughput sequencing and amplicon sequence variants (ASVs) of the ITS1 reads to identify species and some strains as ASVs can distinguish differences as little as one base pair (Callahan et al. 2016). Sequencing yielded an average of approximately 35,550 reads with 131 amplicon sequence variants (ASVs) per sample. In total, there were 6,503 ASVs recovered across the 192 samples excluding sequencing controls. The ITS1 reverse primer was designed to maximize the number of *Pseudo-nitzschia* spp. amplified, and as such was also expected to amplify other genera. Sequences from diatoms outside the *Pseudo-nitzschia* genus and other plankton such as dinoflagellates were recovered in the top 100 most numerous ASVs. Thirty of the top 100 most numerous ASVs had no significant similarity to existing sequences in the National Center for Biotechnology Information (NCBI, Bethesda, MD, USA) nucleotide database and may represent unknown diversity within the marine plankton, or alternatively species lacking ITS1 data.

ASVs were identified as *Pseudo-nitzschia* taxa using a curated database from NCBI sequences (Table S2) and the scikit-learn naïve Bayes machine learning classifier (Pedregosa et al. 2011) which identified 97 ASVs as *Pseudo-nitzschia* at the species level. The number of reads belonging to *Pseudo-nitzschia* spp. in each sample ranged from 19 – 46,314 with an average of 10,655. Nineteen known *Pseudo-nitzschia* species were represented by 53 ASVs. Individual *Pseudo-nitzschia* species ranged from having one to seven ASVs (Table 1). Most taxa represented by two or more ASVs had a dominant ASV found in over double the number of samples compared to the other ASVs of that taxon. However, three taxa had two ASVs which were equally dominant throughout samples: *P. americana, P. calliantha,* and *P. subpacifica.* One group of three similar ASVs were only identified at the genus level for this analysis, and likely represented potential novel diversity of *P. americana* from 95% similar identity via megablast results.

Because of high variability in ITS1 rDNA copy number and unknown copy number per *Pseudo-nitzschia* strain, this study could not equate or transform ITS1 reads to absolute abundance, or even relative abundance, of *Pseudo-nitzschia* species. For example, quantitative PCR has been used before to determine ITS1 copy number in eight *Pseudo-nitzschia* species and a significant variation of two orders of magnitude was found between 16 copies in a *P. delicatissima* isolate to 748 in *P. multiseries* (Hubbard et al. 2014). Thus, in our analyses we used the presence or absence of individual *Pseudo-nitzschia* ITS1 ASVs to circumvent the copy number challenge and examine co-occurring taxa.

### Seasonal cycle of Pseudo-nitzschia spp. diversity

*Pseudo-nitzschia* spp. composition was compared in all samples. Season was the only reliable grouping of ASVs to test for analysis of similarities (ANOSIM) as determined by dispersion of group tests (n = 192, p = 0.771). ANOSIM showed a significant difference between *Pseudo-nitzschia* assemblages during different seasons (p = 0.001, Fig. 5A). Nonmetric multidimensional scaling (NMDS) of the Jaccard distance matrix of the presence and absence of *Pseudo-nitzschia* ASVs showed assemblages were similar across most sites sampled in the same season, including the offshore NES LTER samples (Fig. 5A). Additionally, there was a group of winter NES LTER samples which did not group with other NBay samples (Fig. 5A). Adjacent seasons shared taxa as shown by overlapping points, and there was a general pattern across an annual trajectory connecting adjacent seasons (Fig. 5A). Exceptions to this general seasonal trend were spring closure samples from 2017 that grouped with fall samples from 2018 and 2019 (Fig. 5A).

**Fig 5.**
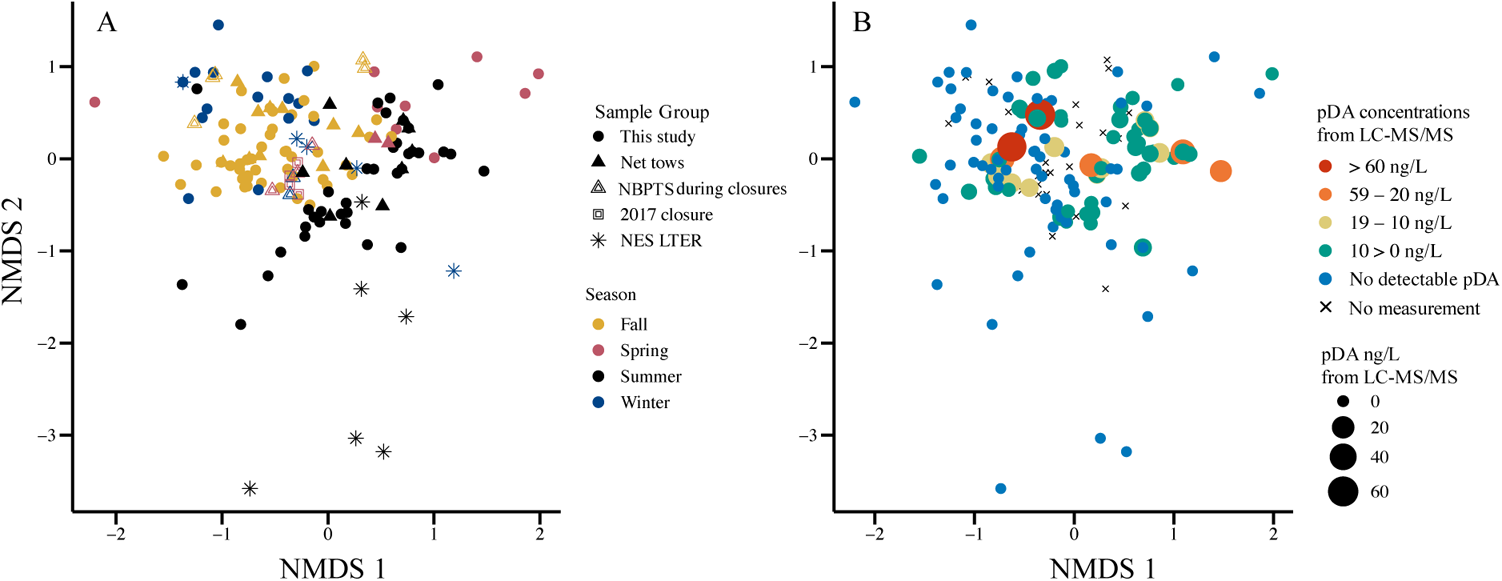
Nonmetric multidimensional scaling (NMDS) of Jaccard distance matrix of the presence or absence of *Pseudo-nitzschia* spp. amplicon sequence variants in samples from Narragansett Bay (NBay), RI and offshore (n = 192). Both plots are the same NMDS with stress = 0.0924. Solution was not reached by defaults, so iterations raised to 100 and dimensions were raised to three but visualized in two dimensions. Views of the same plots in 3-D are available in Fig S8. **[A]** *Pseudo-nitzschia* spp. assemblages across seasons (color) by sample group (shape), including the NBay Long-Term Plankton Time Series (NBPTS) and Northeast Shelf Long-Term Ecological Research (NES LTER) offshore cruises, which has different sample types and sample years. **[B]** Assemblages of *Pseudo-nitzschia* taxa overlaid with concentration of particulate domoic acid (pDA) measured in filtered seawater L^-1^ from LC-MS/MS measurements. Samples where no measurements were taken for LC-MS/MS are indicated with an x, including net tow, precautionary closure, closure, and some NES LTER samples.

The *Pseudo-nitzschia* assemblages co-occurring with elevated pDA concentrations (> 10 ng pDA L^-1^) in the late spring/early summer and late summer/early fall were distinct from each other (Fig. 5). However, there was some overlap of taxa, such as *P. multiseries,* throughout seasons (Fig. 5, Fig. 6). Interestingly, within a season, similar assemblages both do and do not produce DA (Fig. 5). From the BIOENV analysis of the environmental variables of temperature, salinity, DSi, DIP, ammonium, nitrate, nitrite, DIN:DIP, and DIN:Si, it was determined that temperature, DIP, nitrate, and DIN:DIP best correlated with the *Pseudo-nitzschia* ASV dataset (n = 127 samples, correlation = 0.44).

**Fig 6.**
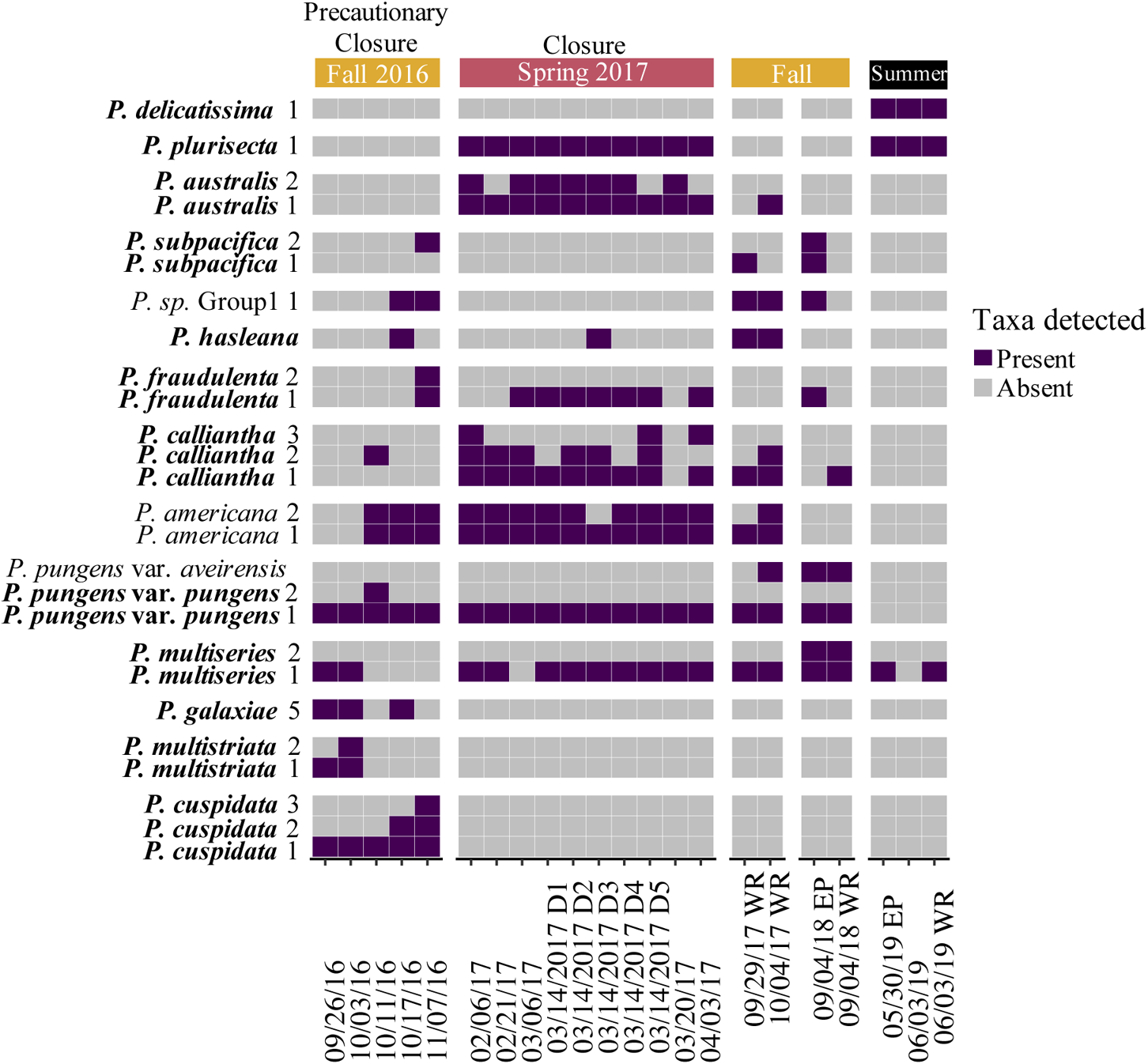
*Pseudo-nitzschia* spp. detected in samples around the 2016 precautionary closure, around the 2017 closure, and from the seven most toxic samples during the study period in Narragansett Bay (NBay), RI. The samples from the 2016 precautionary closure (10/07/16 – 10/29/16) and 2017 closure (02/26/17 – 03/24/17) included those collected at the NBay Long-Term Plankton Time Series (NBPTS) site and RI DEM HAB monitoring sites (D1 – D5). Samples with the highest particulate domoic acid (pDA) concentrations were from the study period from September 2017 – November 2019 at NBPTS, Whale Rock (WR), and East Passage (EP). Samples without a location indicated are from the NBPTS site. Distinct amplicon sequence variants (ASVs) identified as the same species were kept separate. There was a group of ASVs in the data set which could only be identified to the genus level, labeled as “*P. sp.* Group1”. Purple shading indicates ASVs which occurred at >1% relative abundance per sample as present and grey as those not present at < 1% relative abundance or absent in an individual sample. Species known to produce domoic acid (DA) according to Bates *et al*. 2018 are denoted in bold. Corresponding pDA concentrations are 60, 34, 66, 29, 26, 22, and 35 ng pDA L^-1^ seawater filtered for 09/29/17 at WR, 10/04/17 at WR, 09/04/18 at EP, 09/04/18 at WR, 05/30/19 at EP, 06/03/19 at NBPTS, and 06/03/19 at WR, respectively.

*Pseudo-nitzschia* species richness followed a seasonal cycle (Fig. S6). From Levene’s test, it was determined that the number of ASVs by seasonal groupings displayed homogeneity of variances (p > 0.05), following the assumption appropriate for a one-way ANOVA. There was a significant difference in the number of ASVs recovered from samples by season as determined by a one-way ANOVA (p < 0.001), and it was determined from Tukey multiple pair-wise comparisons that fall compared to the other seasons had significant differences in the number of ASVs (p < 0.01). Samples with the highest number of ASVs occurred from August – October, and diversity peaked again in March and decreased in December – January and May (Fig. S7).

The number of *Pseudo-nitzschia* ASVs in NBay samples ranged from 1 – 13 with an average of six ASVs present. The average number of ASVs recovered from the 2016 precautionary closure and 2017 closure samples was higher at eight ASVs. There was no difference between diversity measures when cell counts were under or over the action threshold or when pDA was detected or not.

### Multi-species assemblages of Pseudo-nitzschia spp

Fourteen known DA-producing taxa were found in NBay (Table 1) (Bates et al. 2018). Some of these toxigenic *Pseudo-nitzschia* spp. (*P. multiseries, P. calliantha,* and *P. pungens* var. *pungens*) were frequent and found in 70% or more of the total 2017 – 2019 NBay samples (Table 1). The non-toxic *P. americana* (Bates et al. 2018) was also found in approximately 70% or more of the total 2017 – 2019 NBay samples. The 2016 precautionary closure samples had a lower diversity of toxigenic *Pseudo-nitzschia* (*P. pungens* var. *pungens* and *P. cuspidata)* compared to the higher diversity found in all 2017 closure samples (*P. pungens* var. *pungens*, *P. multiseries, P. americana, P. calliantha, P. fraudulenta, P. plurisecta,* and *P. australis*) (Fig. 6). This study was also the first documentation of eight taxa along the U.S. Atlantic coast (Table 1). Of these newly documented species, *P. hasleana, P. galaxiae,* and *P. multistriata* were present in the closure samples from 2016 (Fig. 6) and from 2017 – 2019, *P. pungens* var. *aveirensis* was present in the fall and co-occurred with *P. pungens* var. *pungens* (Fig. S5). The other newly observed taxa were rarely observed (Fig. S5).

The presence of rare, but highly toxigenic *Pseudo-nitzschia* spp. may augment pDA in NBay. Specifically, *P. australis* is one of the most toxigenic *Pseudo-nitzschia* species (Trainer et al. 2012), and it was present in every 2017 closure sample (Fig. 6) and the NES LTER samples with detected pDA (Fig. S8). In contrast, *P. australis* was absent in the 2016 precautionary closure and infrequent in the 2017 – 2019 NBay samples and the other NES LTER samples (Fig. 6, Fig. S5, Fig. S6, Fig. S8). The 2016 precautionary closure also contained ASVs from *P. multistriata, P. cuspidata,* and *P. galaxiae* that were rarely found in the 2017 – 2019 NBay samples (Fig. 6, Fig. S5).

## Discussion

### Patterns of pDA repeat seasonally under closure thresholds

In NBay, DA-producing *Pseudo-nitzschia* spp. are a newly emergent HAB problem. In this study, a combination of high frequency sampling at several locations, sensitive quantitative pDA measurements and new species delineation methods advanced our understanding of the ecological dynamics of *Pseudo-nitzschia* spp. and toxin production in NBay. The rapid deployment of LC-MS/MS measurements allowed for near real-time detection of pDA as well as the ability to adaptively sample additional sites during occurrences of increasing pDA. The LC-MS/MS method is over two orders of magnitude more sensitive (LOQ = 1 ng pDA L^-1^ seawater filtered) than the Scotia Rapid Tests employed by RI HAB monitoring for the detection of DA in the plankton (LOD = 300 ng pDA L^-1^ filtered seawater (Jellet et al. 2006)). Thus, the repeating seasonal peaks of pDA revealed in this study, even at their highest, would not have been detected by routine RI DEM HAB monitoring. The advantage of this LC-MS/MS method for uncovering HAB dynamics was illustrated during spring 2019. From November 12, 2018 until May 20, 2019, pDA was consistently undetectable at all sites (Fig. 3C). Beginning May 20, 2019, low pDA was detected at the mouth of NBay. Sampling frequency was increased, and the following week (May 28, 2019), pDA levels were 10-fold higher at the NBay entrances (Fig. 3C). There was continued detection of pDA throughout June 2019, and we captured the entire event of pDA increasing, reaching its maximum, and decreasing back to undetectable levels (Fig. 3C).

Our time series sampling from 2017-2019 showed peaks in pDA that increased repeatedly in the fall and summer, generally for two to three weeks (Fig. 3C). These data, along with the 2016 fall precautionary closure, indicate the times of the year when DA production is elevated and potentially a problem in NBay. These patterns would have been missed by solely relying on enumeration of *Pseudo-nitzschia* spp. cells alone, which were much lower than RI DEM action thresholds during the 2017-2019 time series (cf. Fig. 3B, Fig. 3C, Fig. S3). The frequent and highly sensitive pDA measurements allowed us to better understand toxin dynamics that would have been masked by using elevated cell counts as a proxy. The highest pDA concentration (66 ng L^-1^) measured by our study was similar to pDA levels in the Gulf of Maine in 2012 (60 ng L^-1^), but much lower than in the 2016 Gulf of Maine *P. australis* bloom (37.5 µg L^-1^) (Clark et al. 2019). Even though the pDA detected during our study period was below closure concerns, our data show that DA is seasonally present, which could lead to bioaccumulation in the food web.

The impacts of repetitive non-acute DA doses on humans is currently unknown; however, low-level DA consumption has been associated with everyday memory problems in humans (Grattan et al. 2018), decreased memory in infants through prenatal ingestion in nonhuman primates (Grant et al. 2019), and spatial learning impairment and hyperactivity in mice (Lefebvre et al. 2017). Further investigation of possible acute and chronic DA vectors in NBay is necessary. Vector species including oysters, scallops, squid, and carnivorous snails are known to bioaccumulate DA, but little is known about their bioaccumulation dynamics such as species-specific toxin purging rates and human diet frequency in NBay (Trainer et al. 2012; Bates et al. 2018).

### Season and temperature structure Pseudo-nitzschia spp. assemblages

When we initiated our study in 2017, we hypothesized that the DA closures in NBay were either caused by the entrainment of a particularly toxic species typically not present at high abundances in NBay or by changing environmental conditions augmenting toxin production in resident *Pseudo-nitzschia* spp. assemblages. We have demonstrated support for both hypotheses. The new ITS1 high-throughput sequencing methods deployed in this study both uncovered new *Pseudo-nitzschia* species diversity and identified strains of the same species. We identified species not previously known in NBay, as well as eight species previously unreported along the U.S. Atlantic coast (Table 1). New *Pseudo-nitzschia* spp. diversity uncovered by using high-throughput sequencing is similar to findings from another NBay molecular study that demonstrated newly observed species of the diatom genus *Thalassiosira* (Rynearson et al. 2020). The application of high-throughput sequencing with the ITS1 instead of with ARISA increased the ability to delineate certain species that have the same ITS fragment length such as *P. australis* and *P. seriata* through the sequence identity as well as additional *Pseudo-nitzschia* species (e.g. *P. calliantha, P. pungens* var. *aveirensis,* and *P. hasleana*) from the more inclusive primer pair used in this study (Hubbard et al. 2008).

Several co-occurring species were found frequently in more than 70% of samples from 2017-2019. We considered these common residents of NBay: *P. multiseries, P. pungens* var. *pungens,* and *P. calliantha* (Table 1). Notably, these three species were also present during the 2016 precautionary closure (Fig. 6). Our study also showed that recurring resident *Pseudo-nitzschia* assemblages in Narragansett Bay can be driven to toxin production. Distinct multi-species assemblages of *Pseudo-nitzschia* grouped by season and recur interannually (Fig. 5). Assemblages sampled within a season had different pDA concentrations despite containing similar species (Fig. 5).

*Pseudo-nitzschia* spp. have been observed to persist across the annual range of temperatures experienced in NBay (Miller and Kamykou 1986; Dortch et al. 1997; Delegrange et al. 2018). During the 2017-2019 sampling period, NBay surface water temperature ranged from - 1.1 to 25.5 °C, and *Pseudo-nitzschia* spp. were observed year-round. Low temperatures may be controlling the mostly non-toxic winter assemblage, which included *P. americana* (Fig. 4C, Fig. S5). However, pDA in NBay was detected across temperatures ranging from 3.1 - 24°C. Temperature is likely a major driving force of the seasonal species groupings, and has been shown to influence other diatom assemblages in NBay (Karentz and Smayda 1984; Canesi and Rynearson 2016; Rynearson et al. 2020). It is notable that the water temperatures were very different during the 2016 and 2017 closures with 15 – 20 °C in October 2016 and 1 - 5 °C in March 2017 (https://web.uri.edu/gso/research/plankton/) and each contained different species assemblages (Fig. 5, Fig. 6). These differences demonstrate that species assemblages forming under different NBay temperatures matter more for DA production potential than does temperature itself as a direct driver of DA production.

### Nitrogen availability may vary pDA production by similar Pseudo-nitzschia spp. assemblages

The repeating summer and fall pDA peaks correlated with low DIN concentrations (Fig. 3, Fig. 4, Fig S4). Therefore, nutrient dynamics likely enhance toxin production in toxin-capable assemblages in NBay. There have been other observations of increased DA production under nitrogen-limiting conditions, depending on species, cell cycle stage, and nitrogen substrate (Trainer et al. 2012). For example, strains of *P. multiseries* produce DA under nitrogen-limiting conditions with a variety of nitrogen substrates (Hagström et al. 2011; Trainer et al. 2012). In NBay, *P. multiseries* was a common member of both the toxic summer and fall assemblages that occurred during periods of low DIN (Fig. S3, Fig. 3D). It seems counter-intuitive to consider that nitrogen stress would lead to elevated DA production in *Pseudo-nitzschia* spp. as nitrogen is a requirement for the production of DA. Glutamine, which results from nitrogen assimilation, is a precursor in the pathway elucidated for DA biosynthesis in *P. multiseries* (Brunson et al. 2018). However, studies on the remodeling of nitrogen metabolism under nitrogen stress in the non-DA producing diatom *Phaeodactylum tricornutum* have shown that enzymes in glutamine biosynthesis are upregulated by nitrogen limitation (Levitan et al. 2015). Additionally, a switch from DIN to urea as a nitrogen source may support *Pseudo-nitzschia* growth and increased toxicity as well (Howard et al. 2007; Kudela et al. 2008). Currently it is unknown how urea, an anthropogenic nitrogen source, impacts DA production in NBay, which may be useful to examine in future studies.

In addition to a relationship between low DIN and elevated pDA, there may be multiple co-occurring ecological factors eliciting species-specific responses of cell growth and toxin production. Notably, strains of the same *Pseudo-nitzschia* spp. may respond differently (Thessen et al. 2009; Sahraoui et al. 2011; Markina and Aizdaicher 2016). In NBay, strain-specific responses may play a key role in toxin production, as most species had more than two ASVs assigned (Table 1). For example, the fall species assemblages produced the highest pDA during our 2017 - 2019 study period. This could be attributed to toxin production by unique species constituents distinct from the spring assemblages, or other conditions in addition to low DIN forcing greater toxin production in species common to both fall and spring such as *P. multiseries* or *P. pungens* var. *pungens* (Fig. S5). Complicating the challenge of deciphering ecological factors contributing to DA production is the long-standing mystery of why some *Pseudo-nitzschia* spp. produce DA and whether DA production confers an evolutionarily or ecological benefit (Zabaglo et al. 2016). Other biotic factors unexamined in this study could also influence toxin production such as bacterial communities (Bates et al. 1995; Sison-Mangus et al. 2016) and grazing pressure (Bates et al. 2018).

### Pseudo-nitzschia australis as a species of concern and management implications

In NBay, the absence of DA closures prior to 2016 - 2017 was likely not due to a lack of monitoring or under sampling. RI HAB monitoring involves three levels: an action threshold for *Pseudo-nitzschia* spp. cell counts, the Scotia Rapid Test for pDA in the plankton, and a management threshold for DA in shellfish meat. Sampling is expansive with 38 sites ranging from offshore Block Island, RI to coastal ponds to upper NBay. This effort totals over 300 discrete samples annually, which are concentrated from 20 L of surface seawater across a 20 μm plankton net (2020).

In October 2016, the first HAB-biotoxin shellfish harvest closure due to elevated plankton-associated DA lasted for 26 days in NBay (Fig. S1A). It was a precautionary closure since DA in the shellfish meat did not exceed the management threshold; however, DA was detected in wild shellfish meat, including quahogs (*Mercenaria mercenaria*) and mussels (*Mytilus edulis*). During the 2016 closure period, nine toxigenic species were found at the NBPTS site, but notably, *P. australis* was absent (Fig. 6). Samples from nearby Massachusetts during the same time period on October 11, 2016 included multi-species assemblages with *P. pungens* var. *pungens* and others, as well as an absence of *P. australis*. Surprisingly, this differs from *P. australis* present in Maine and Canada during their coinciding 2016 closures (Bates et al. 2018; Clark et al. 2019).

In 2017, a second closure in NBay occurred when DA was detected in shellfish meats collected near the NBay entrances adjacent to RI coastal waters, and *P. australis,* which may have originated from offshore populations, was present in all samples sequenced (Fig. S1B, Fig. 6). The presence of *P. australis,* with possible additional toxin contributions from *P. multiseries, P. plurisecta, P. fraudulenta, P. calliantha,* and *P. pungens* var. *pungens,* likely was responsible for the 2017 closure. From 2017-2019, *P. australis* was not commonly observed, except for fall 2017, winter and spring 2018, and fall 2019 when pDA was less than 10 ng L^-1^ (Fig. S5). The winter 2018 samples coincided with offshore NES LTER sampling, where *P. australis* was present at all stations and the station closest to NBay had 15 ng pDA L^-1^ (Fig. S8A). Altogether, these data suggest that *P. australis* is not a resident species of NBay, but instead likely introduced from offshore. This taxon may be particularly problematic in regard to DA production in NBay. Perhaps in 2016, the geographic barrier of Cape Cod or differing water masses separated *Pseudo-nitzschia* spp. assemblages, up and down the Atlantic Northeast. Therefore, one assemblage contained *P. australis* in Maine and Canada and there was a different toxigenic assemblage in RI and Massachusetts, which led to two seemingly unrelated regional closures. Differences in water masses have been shown to separate *Pseudo-nitzschia* species assemblages in other environments (Bowers et al. 2018; Clark et al. 2019). Also, this trend of an expanded geographic range for *P. australis* may be part of a larger pattern in the Atlantic Ocean: *P. australis* expanded into northern European waters (Trainer et al. 2012) and was not documented until 1994 in northwest Spain (Míguez and Fernlindez 1996) or until 1999 in Scottish waters (Gallacher et al. 2001).

The introduction of *P. australis* into NBay may have been driven by climate change augmented by the decreased nitrogen loading which may have allowed the cells to persist in NBay. Along the US Pacific Coast, increased *P. australis* growth rates were linked to climate change-related warm water anomalies and low nutrients from a stratified euphotic zone (McCabe et al. 2016; Ryan et al. 2017). *P. australis* persisted in the nutrient-depleted stratified waters and achieved high growth rates when nutrients were resupplied from upwelling (McCabe et al. 2016; Ryan et al. 2017; McKibben et al. 2017). Notably, in NBay, nitrogen loading declined significantly from recent implementation of tertiary sewage treatment (Oviatt et al. 2017; Oczkowski et al. 2018), and waters have warmed by 1.4 – 1.6 °C since 1960 (Fulweiler et al. 2015) as well as the offshore waters on the North Atlantic continental shelf by 0.37 °C (Chen et al. 2020). This combination of altered nutrient status and background of elevated temperature may contribute to conditions augmenting *Pseudo-nitzschia* HABs in NBay.

The contrasting dynamics we observed between *Pseudo-nitzschia* originating from offshore vs. resident populations have implications for HAB management in NBay and regionally, as these data show the importance of species-specific dynamics in toxic events. Although no closures occurred from September 2017-November 2019, we established baseline data for seasonal *Pseudo-nitzschia* spp. assemblages that regularly recur, especially during detectable pDA peaks in the fall and summer. The presence of particularly toxigenic species, such as *P. australis*, during these time periods will inform when environmental and species intersections are concerning in regard to HAB formation.

### Conclusions

In this study, we showed a high-resolution time series of pDA concentrations and *Pseudo-nitzschia* spp. assemblages which revealed seasonal patterns in NBay that appear to recur annually. These observations will assist in predicting timing of future DA closures, with the identification of possible abiotic factors, temperature and nitrogen, contributing to toxin production. High-throughput sequencing of the ITS1 rDNA region has provided important multi-species composition data, including identifying rare but potentially highly toxic species (i.e. *P. australis*) and previously unknown taxa to the region along with the composition of resident assemblages that may be driven to higher toxin production. Because the highest pDA concentrations were observed at the NBay mouths, these entrances may serve as sites for sentinel mussels and other tools to monitor the introduction of offshore species. NBay managers and shellfish harvesters must prepare for possible HAB event increases, as this study has shown low levels of pDA present in the fall and summer that reoccur each year and their magnitude may change as it did in 2017 leading to a DA closure. Additional temporal introduction of highly toxigenic species such as *P. australis,* responding to shifts in larger oceanographic patterns and warming water temperatures, may provide increasingly favorable conditions for more severe HABs in NBay and along the Northeast Atlantic.

## Acknowledgements

We acknowledge S. Anderson, K. Bell, D. Fontaine, L. Holland, M. McKenzie and J. Vazquez-Custodio and for sampling assistance; S. Barber (R/V *Cap’n Bert*), H. Vincent II (R/V *Hope Hudner*), and S. Granger and D. Ullman (R/V *Zostera*), and the R/V *Endeavor* crew for field assistance. We thank S. Beaulieu for data management assistance, Z. Pimentel for help with DNA sequencing analysis; J. Rines for the *Pseudo-nitzschia* culture, and C. McManus for helpful conversations. The URI GSO NBPTS is supported by URI and the RI Department of Environmental Management. The RI Genomics and Sequencing Center (DNA sequencing) and the Marine Science Research Facility (Lachat nutrients) are supported by the NSF EPSCoR Cooperative Agreements (EPS-1004057, OIA-1655221). Spectroscopic and spectrometric measurements were made in the URI RI-INBRE Centralized Research Core Facility, supported by the Institutional Development Award (IDeA) Network for Biomedical Research Excellence from the National Institute of General Medical Sciences of the National Institutes of Health (P20GM103430). This research was supported by RI Sea Grant Awards to M.B., B.D.J. (NA18OAR4170094), L.M. (NA14OAR4170082), T.A.R. (NES LTER OCE 1655686) and the NSF RI C-AIM EPSCoR Cooperative Agreement (OIA-1655221). S.V. was supported by the URI GSO NSF REU Program (OCE-1757572).

