## Supplemental Text and Figures for "Emerging harmful algal blooms caused by distinct seasonal assemblages of the toxic diatom *Pseudo-nitzschia* in Narragansett Bay, RI, USA"

**This PDF file includes:**

Supplementary Text

Figures S1 to S8

Tables S1 to S2

SI References

**Supplementary Materials and Methods**

***Pseudo-nitzschia* spp. enumeration.** Between 15 and 50 mL of whole seawater was preserved with 1% acidic Lugol's solution in glass or plastic containers (with a silica bead later added). Samples were stored at 4 °C for less than a year before being counted. One mL of sample was placed in a Sedgewick-Rafter counting chamber (Science First / Wildco, Yulee, FL, USA), *Pseudo-nitzschia* spp. cells were identified at the genus level and counted using 20x phase contrast light microscopy on a BX40 (Olympus America Inc., Melville, NY, USA). *Pseudo-nitzschia* spp. cell counts from the NBPTS were conducted on live samples using the same enumeration protocol on an Eclipse E800 (Nikon Instruments Inc., Melville, NY, USA). Cell counts by same-day NBPTS on the Eclipse E800 were preferentially used over our counts performed in the BX40, which made for the same data shown in from September 2017 – November 2019 for the NBPTS site in Fig. 2 and Fig. 3 except for any additional excursions by our team at that site.

**Plankton DNA extraction methods.** DNA was extracted using a modified version of the DNeasy Plant DNA extraction kit (Qiagen, Germantown, MD, USA) with an added bead beating step for 1 minute and QIA-Shredder column (Qiagen, Germantown, MD, USA) as reported in Chappell et al. 2019. Additionally, DNA was eluted in 30 µL with a second elution step of either 30 or 15 µL to maximize DNA yield. DNA was assessed for quality with a Nanodrop spectrophotometer (Thermo Fisher Scientific Inc., Waltham, MA, USA) and quantified using a Qubit fluorometer (Invitrogen, Carlsbad, CA, USA) with the Broad Range dsDNA and High Sensitivity dsDNA kits (Thermo Fisher Scientific Inc., Waltham, MA, USA). DNA yields reported by the Qubit ranged from below the limit of detection to 26.5, with an average of 2.0 ng DNA / µL eluent. NBPTS samples from October 2016 and March 2017 had an average of 300 mL surface seawater passed over a 25 mm 0.2 μm filter, were extracted following existing NBPTS methods of DNA extraction using the DNeasy Blood and Tissue Kit (Qiagen, Germantown, MD, USA) with an added bead beating step (Canesi and Rynearson 2016), and yielded average 0.9 ng DNA / μL eluent as measured by the Qubit. Net tow samples had 50 mL of concentrate passed across a 0.22 μm pore size Sterivex filter unit (MilliporeSigma, Burlington, MA, USA), and were extracted with the same modified DNeasy Plant DNA extraction protocol as above, with 4x volumes of AP1 buffer and RNase A and beads added to the unit to account for the larger sample surface area, extraction occurring within the capped unit itself to maximize yield, and then the lysate removed with a sterile syringe and subsequent steps with adjusted volumes as appropriate. As expected, DNA yields were higher from the Sterivex units ranging from 2.4 – 54.0 ng DNA / μL eluent with an average of 13.7 ng DNA/ μL elution as measured by the Qubit. For the March 13, 2017 NBay samples, 125 mL of surface seawater was passed across a HV filter and extracted with the DNeasy Plant DNA extraction kit with scissors and no beads. As measured by the Qubit, the average DNA yield was 3.7 ng DNA / μL eluent.

**Controls for Sequencing.** A negative control sample was prepared from a blank 25 mm 5.0 μm polyester membrane filter using extraction reagents and had no detectable DNA using the Qubit fluorometer (Invitrogen, Carlsbad, CA, USA). Positive controls were constructed from mock communities comprised of two known *Pseudo-nitzschia* species from monocultures: *P. subcurvata* collected from the Southern Ocean and *P. pungens* var. *pungens* isolated from NBay (provided by J. Rines). One positive control contained equal concentrations of 1 ng extracted DNA. The second positive control was created from equal cell abundances of cultures prior to extraction. These negative and positive controls were amplified and sequenced on the same plate as the other environmental samples.

***Pseudo-nitzschia* spp. ITS1 PCR methods.** Primers (Integrated DNA Technologies, Coralville, IA, USA) were HPLC purified, resuspended in 1x Tris-Acetate-EDTA (TAE) buffer, and then working stocks created in diethylpyrocarbonate (DEPC)-treated $H_2O$. About 4 ng of extracted DNA was used for each PCR reaction. If, according to the Qubit quantification, the DNA concentration was less than 2 ng $\mu L^{-1}$ or below the limit of detection, it was then used as is, and just 2 μL was added to the PCR reaction. PCR reactions were set up on ice, in a 1x reaction in 25 μL total volume. Final primer concentration was 0.5 μM and polymerase was Phusion Hot Start High-Fidelity Master Mix (Thermo Fisher Scientific Inc., Waltham, MA, USA). There were two cycles with different annealing temperatures, the first with an annealing temperature specific to the loci-specific region and the second set of cycles with an annealing temperature that also takes the MiSeq adapter sequence into account (Canesi and Rynearson 2016). PCR conditions used were initial denaturation for 30 seconds at 98 °C, 15 cycles of the following: denaturation for 10 seconds at 98 °C, annealing for 30 seconds at 64.1 °C , extension for 30 seconds at 72 °C, and 15 cycles with the same conditions except a higher annealing temperature of 72 °C, and then a final extension for 10 minutes at 72 °C , and a holding temperature of 10 °C until stored in the -20 °C freezer. PCR products were visualized on a 1% agarose gel before submission to the RI Genomics and Sequencing Center (Kington, RI, USA).

**Taxonomic determination of *Pseudo-nitzschia* ASVs using BLAST.** All of the 6,503 ASVs recovered from the 192 environmental samples were run through a megablast search using BLAST+ v2.9 with the nt database downloaded on October 4, 2020. In addition to the 97 ASVs identified as a specific *Pseudo-nitzschia* species from the QIIME2 pipeline, there were 115

ASVs identified as a *Pseudo-nitzschia* taxon with greater than 75% query coverage and these were manually examined. It was determined by visual inspection that the 11 ASVs identified as *P. pungens* PC50 were likely *Cylindrotheca* instead, and the 85 ASVs which were closest to the *P. delicatissima* KJ22-0.2-69 environmental clone was most closely related to a known *Nitzschia* isolate sequence from subsequent BLAST searches. This left 19 ASVs of interest. Nine have >98% query coverage and >98% identity with known *Pseudo-nitzschia* sequences, so they are referred to as the specific *Pseudo-nitzschia* species match. The additional 10 ASVs were identified as the genus and grouped by similarity with each other. These genus level ASVs have < 96% identity to existing sequences in the database. In total, there were 113 ASVs from the 192 samples that appeared to be of reliable *Pseudo-nitzschia* origin.

**Mass Spectrometry methods for DA analysis.** Samples were chromatographically separated on a Kinetex C18 column (150 mm x 4.6 mm, 2.6 µm, Phenomenex, Torrance, CA, USA) at a flow rate of 0.4 mL $min^{-1}$. Mobile phase solvents were $H_2O$ (A) and MeOH (B) modified with 0.05% formic acid and used in a gradient: for 5 minutes, 95% A and 5% B were held with the eluent sent to waste for the first 2 min, then from 5 to 15 min the % of B increased to 50%, followed by a final change to initial conditions (95% A and 5% B) from 16 to 20 min. The peak of DA eluted at 11.00 min. LC-MS/MS with MRM was employed for sensitivity and selectivity in DA detection and quantification. Analysis was carried out in positive mode, and three transitions from the protonated DA molecule were used: $m/z$ 312 → 266, $m/z$ 312 → 248, and $m/z$ 312 → 193.

**DA analysis from mussel tissue.** The mussels were frozen (-80 °C) until extraction. Following a modified protocol of the Scotia Rapid Test for bivalve tissue, mussels were thawed to room temperature, opened with a knife, and briefly rinsed under cool deionized water to remove salt and sand (Protocol available from Scotia Rapid Testing Ltd. in Nova Scotia, Canada: http://www.jellett.ca/pdf/Bivalve_Tissue_Preparation.pdf). Shellfish meat was removed, and its wet weight was recorded. Samples were hand homogenized in individual tubes for 2 min. A 2.5:1, isopropanol:5% acetic acid, solution was added in equal parts to mussel homogenate. Samples were vortexed for 30 s every 10 min for one hour. Samples were then passed through a small Buchner funnel with Grade 1 filter paper (11 μm particle retention; Whatman n.k.a. Cytiva, Marlborough, MA, USA) under vacuum pressure to produce the mussel extract. The solvent was removed under a stream of $N_2$. Chromatography methods were modified for DA analysis (Beach et al. 2015). Briefly, a strong anion exchange column was conditioned with MeOH, followed by milliQ $H_2O$, then 50:50 $H_2O$:MeOH. The extracts were reconstituted in $H_2O$:MeOH and added to the column. The column was washed with 50:50 $H_2O$:MeOH then the DA from the extract was eluted from the column using $CH_3CN$ and 0.5% formic acid. Samples were then subjected to the same LC-MS/MS analysis with MRM as the pDA samples.

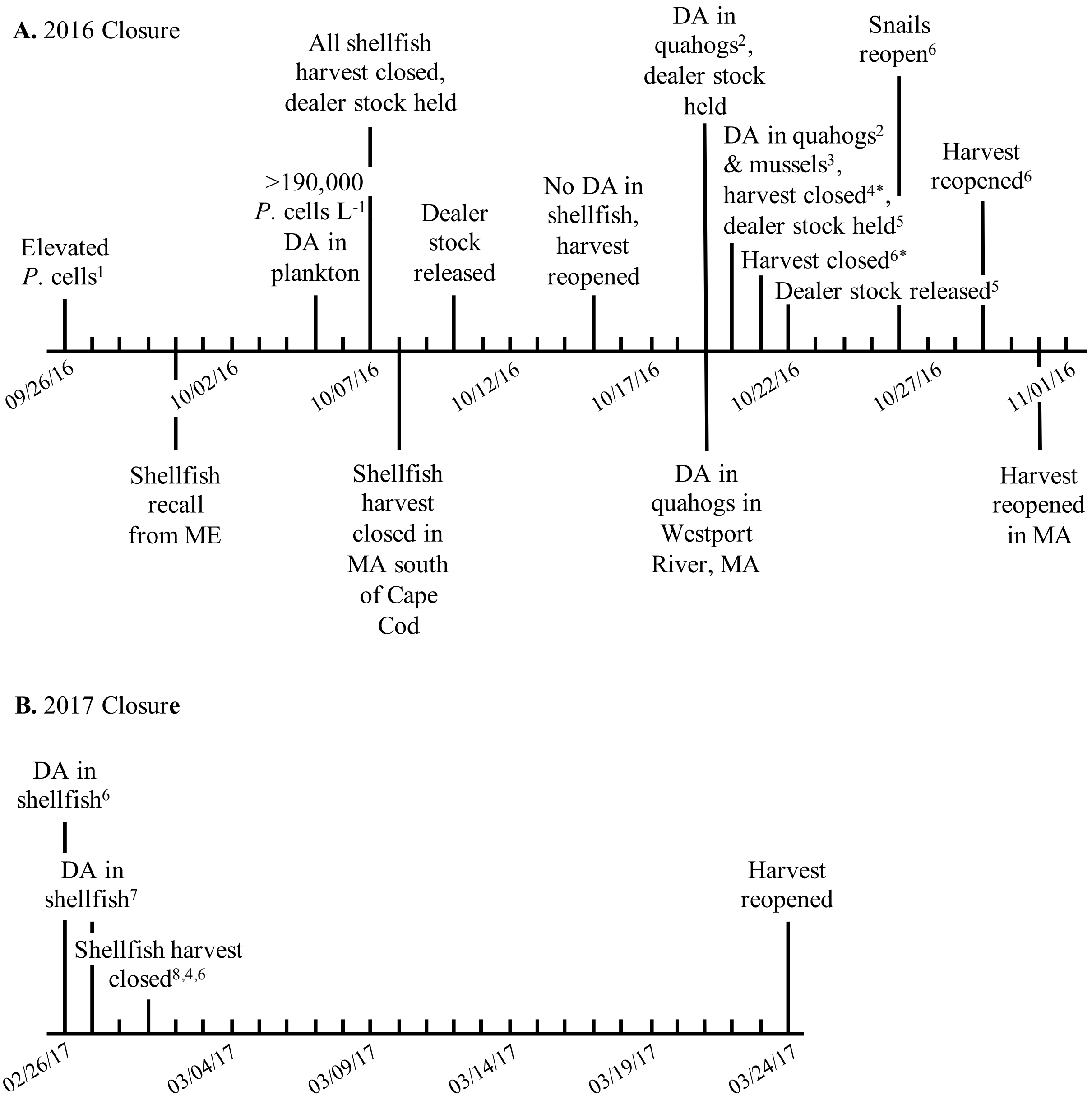


**Fig S1.** Timeline of events related to the **[A]** 2016 precautionary shellfish harvest closure cumulatively from 10/07/16 – 10/29/16 and **[B]** 2017 shellfish harvest closure from 02/26/17 – 03/24/17 in Narragansett Bay, Rhode Island (RI) due to domoic acid, with concurrent 2016 events in Massachusetts (MA) and Maine (ME) underneath. Superscript number specifies location in RI: 1.) Newport Harbor 2.) Sakonnet Harbor 3.) Fort Wetherill 4.) Lower Sakonnet River 5.) East Passage 6.) Lower NBay 7.) Jamestown, Newport, Little Compton 8.) RI Sound. Closures denoted with * indicate that carnivorous snail harvest was prohibited as well. Carnivorous snails include whelk and moon snails.

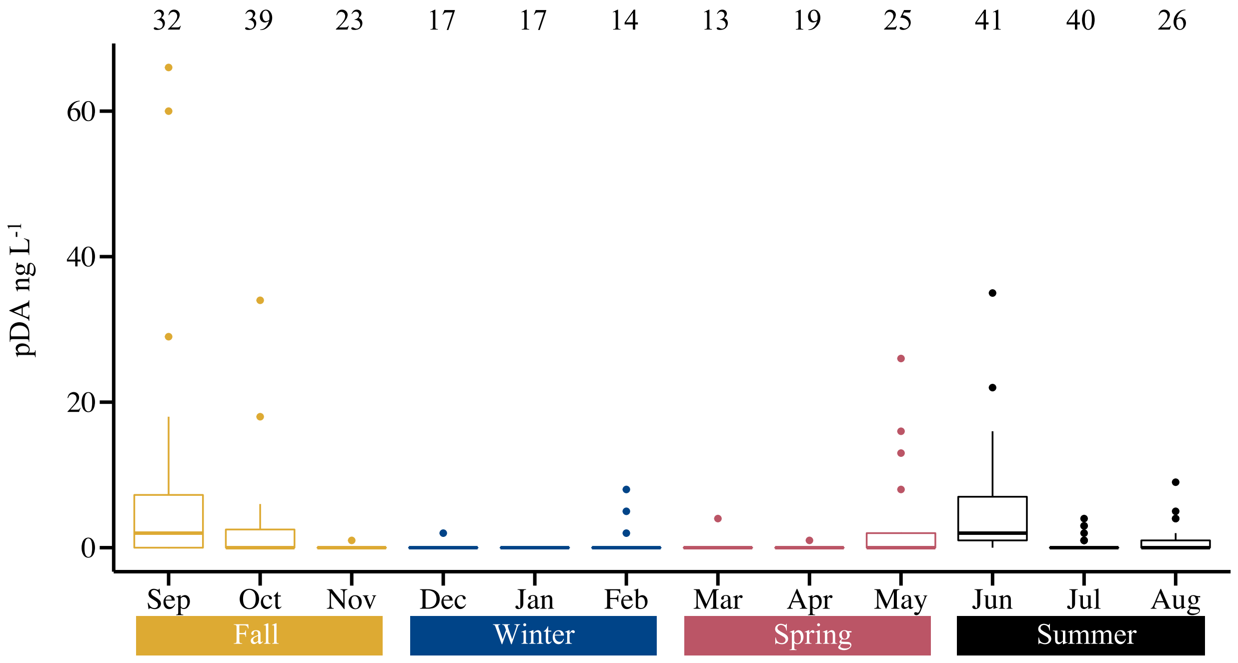


**Fig S2.** Monthly averages of particulate domoic acid (pDA) ng $L^{-1}$ seawater filtered at NBay sites sampled during this study from Sep 2017 – Nov 2019 excluding offshore cruises (n = 306). Months are colored by season, with Sep, Oct, and Nov as Fall; Dec, Jan, and Feb as Winter; Mar, Apr, and May as Spring; and Jun, Jul, and Aug as Summer. Number of samples per month are shown along the top. The 50$^{th}$ percentile is the median, with the whisker shown as 1.5 times the 75$^{th}$ percentile. Outliers are shown as points, which are greater than 1.5 times the interquartile range above the 75$^{th}$ percentile.

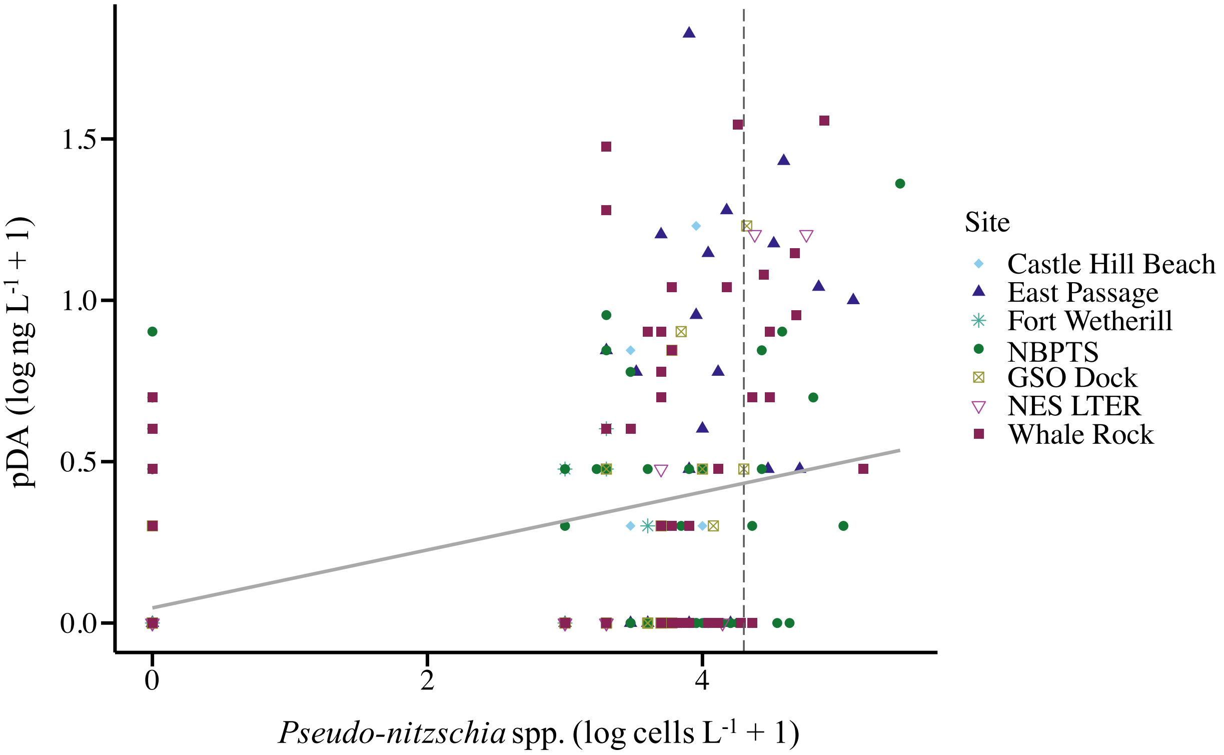


**Fig S3.** Linear relationship between *Pseudo-nitzschia* spp. cell count abundance in surface seawater and concentration of particulate domoic acid (pDA) measured in the same sample when considering all NBay sites and NES LTER samples (n = 305, y = 0.09x + 0.05, $R^2$ = 0.19, p = 8.08 x $10^{-16}$). Values are $Log_{10}$-transformed, with +1 to all values to create non-zero values in the transformation. The vertical dotted line is log-transformed action threshold for RI HAB monitoring of 20,000 *Pseudo-nitzschia* cells $L^{-1}$.

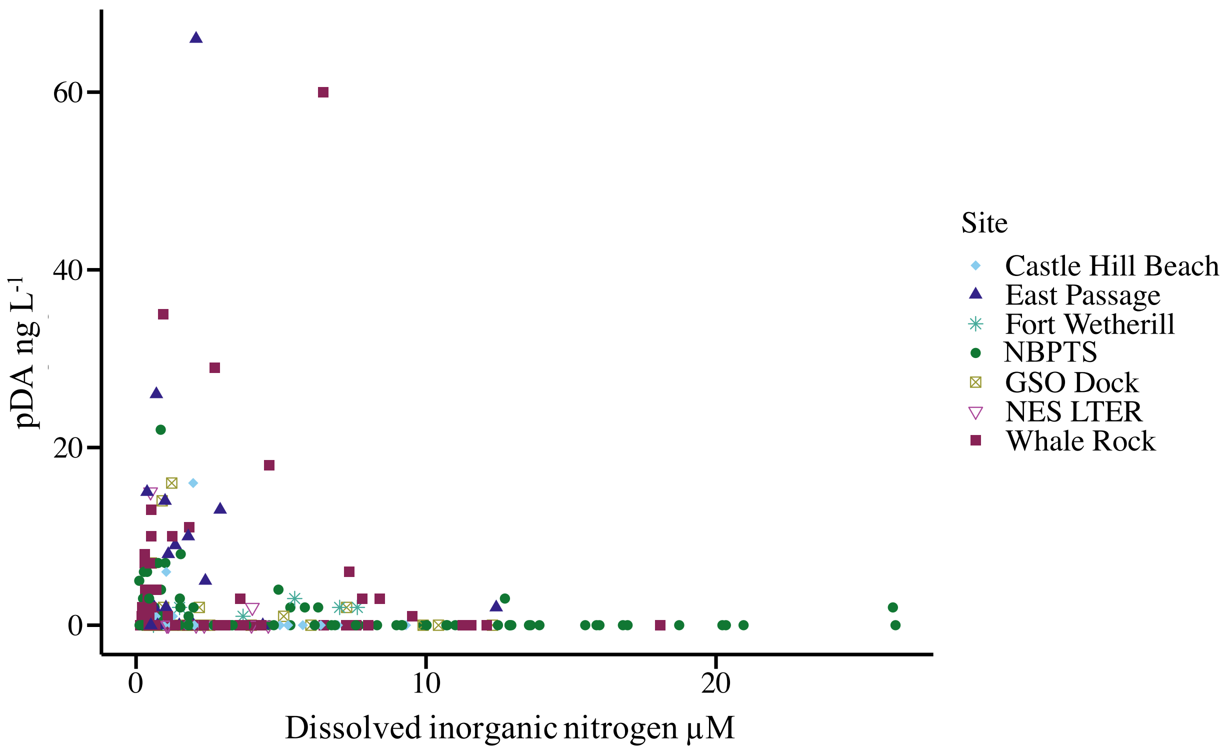


**Fig S4.** Scatterplot of particulate domoic acid (pDA) concentrations in ng $L^{-1}$ from surface seawater samples and dissolved inorganic nitrogen (DIN) µM measurements, including NBay sites and NES LTER samples (n = 294).

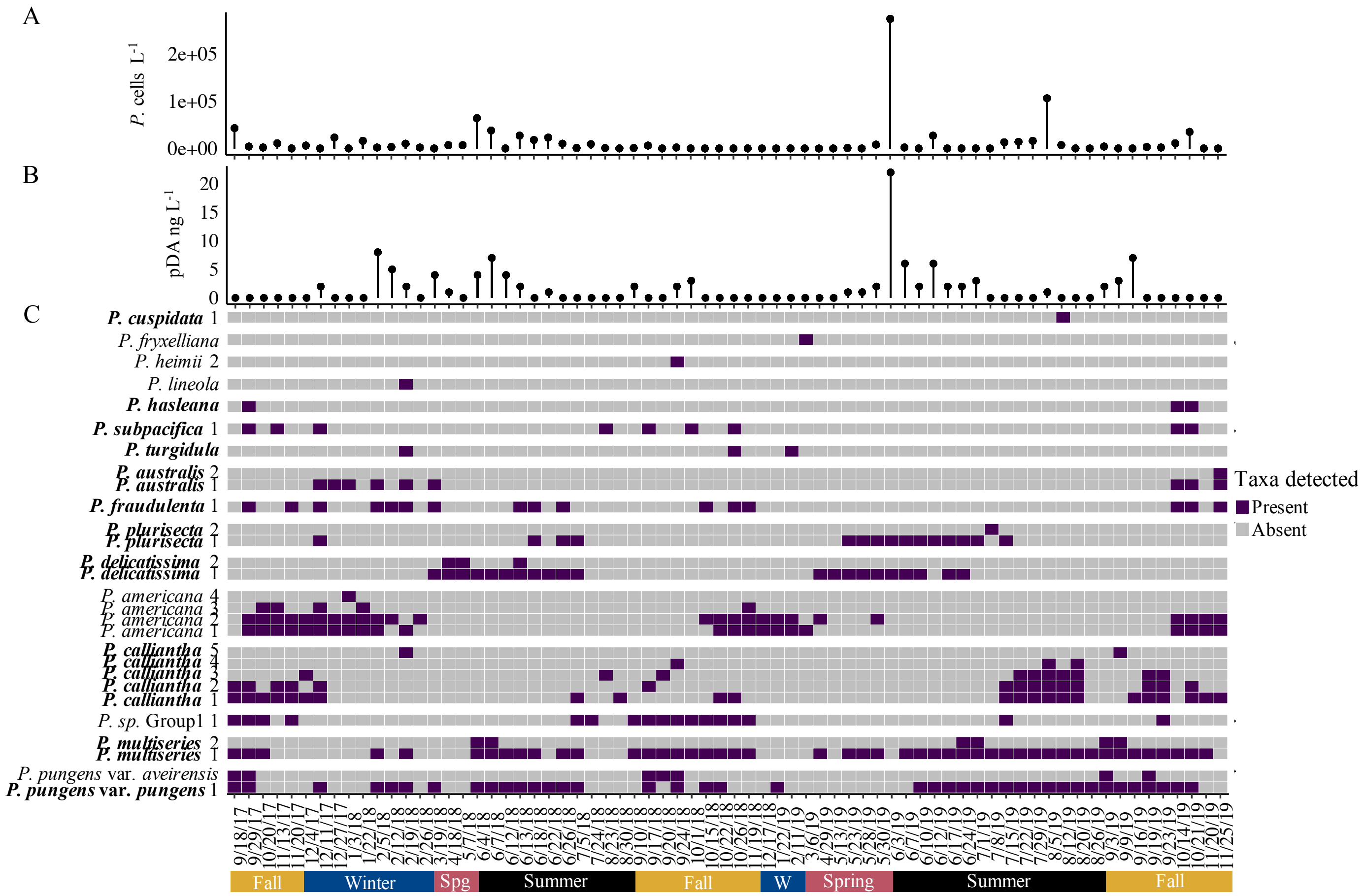


**Fig S5.** Patterns of *Pseudo-nitzschia* spp. identification, cell abundance, and particulate domoic acid (pDA) at corresponding samples at the Long-Term Plankton Time Series (NBPTS) during the study period from Sept 2017 – Nov 2019 (n = 70). Season is indicated along the bottom with "Spg" as Spring and "W" as Winter. **[A]** Cell abundance of *Pseudo-nitzschia* spp. in surface whole seawater $L^{-1}$. **[B]** pDA measured in filtered surface seawater $L^{-1}$. **[C]** Presence or absence of *Pseudo-nitzschia* amplicon sequence variants (ASVs) recovered from high-throughput sequencing of the ITS1 region. Distinct ASVs identified as the same species were kept separate. There was a group of ASVs in the data set which could only be identified to the genus level, labeled as "*P.* sp. Group1". Purple shading indicates ASVs which occurred at >1% relative abundance per sample as present, and grey as those not present at < 1% relative abundance or absent in an individual sample. Species known to produce domoic acid (DA) according to Bates *et al.* 2018 are denoted in bold.

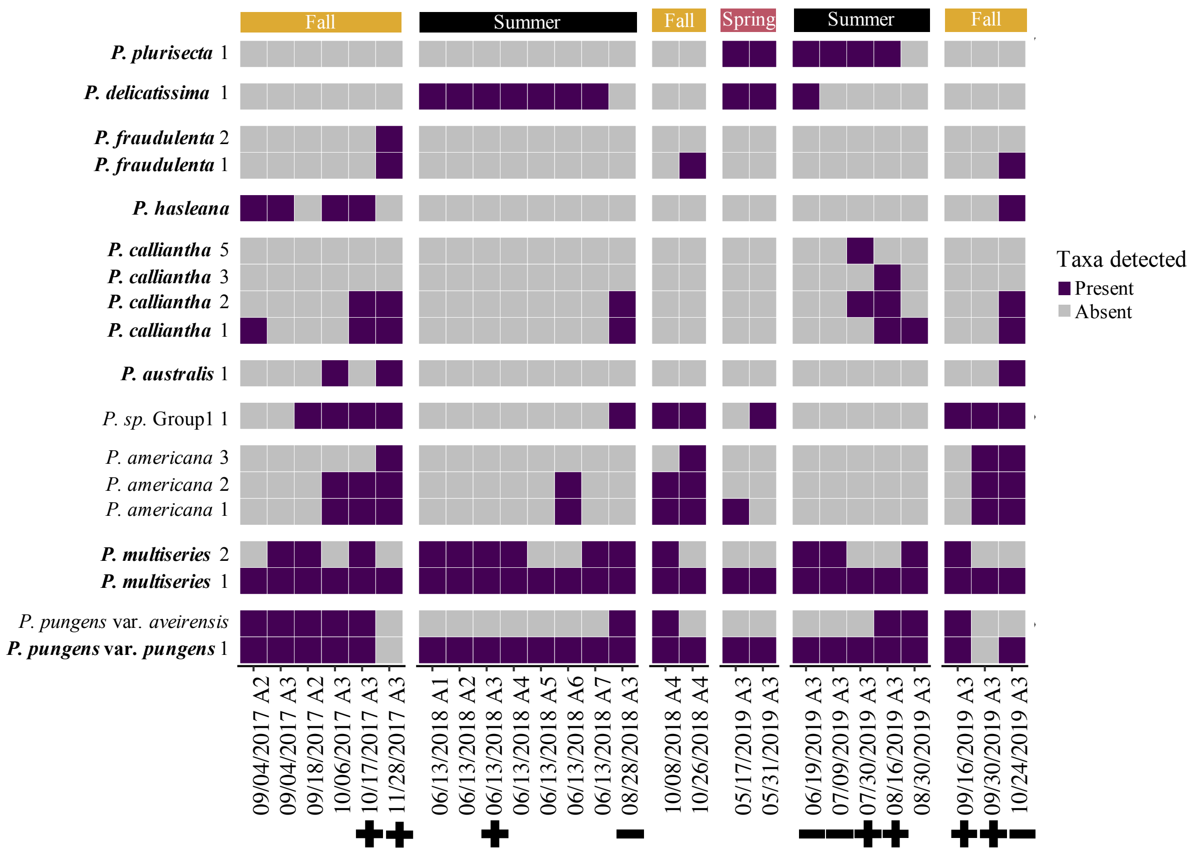


**Fig S6.** *Pseudo-nitzschia* ASVs and their taxonomic identity detected from Narragansett Bay, RI net tow samples (n = 26). Sample date and location (A#) are shown along the bottom, with season along the top. Underneath the date are the results of Scotia Rapid Tests for the samples: plus sign for positive result, minus sign for negative result, and blank if no test was performed. Distinct ASVs identified as the same species were kept separate. There was a group of ASVs in the data set which could only be identified to the genus level, labeled as "*P.* sp. Group1". Purple shading indicates ASVs which occurred at >1% relative abundance per sample as present and grey as those present at < 1% relative abundance or absent in an individual sample. Species known to produce domoic acid (DA) according to Bates *et al.* 2018 are denoted in bold.

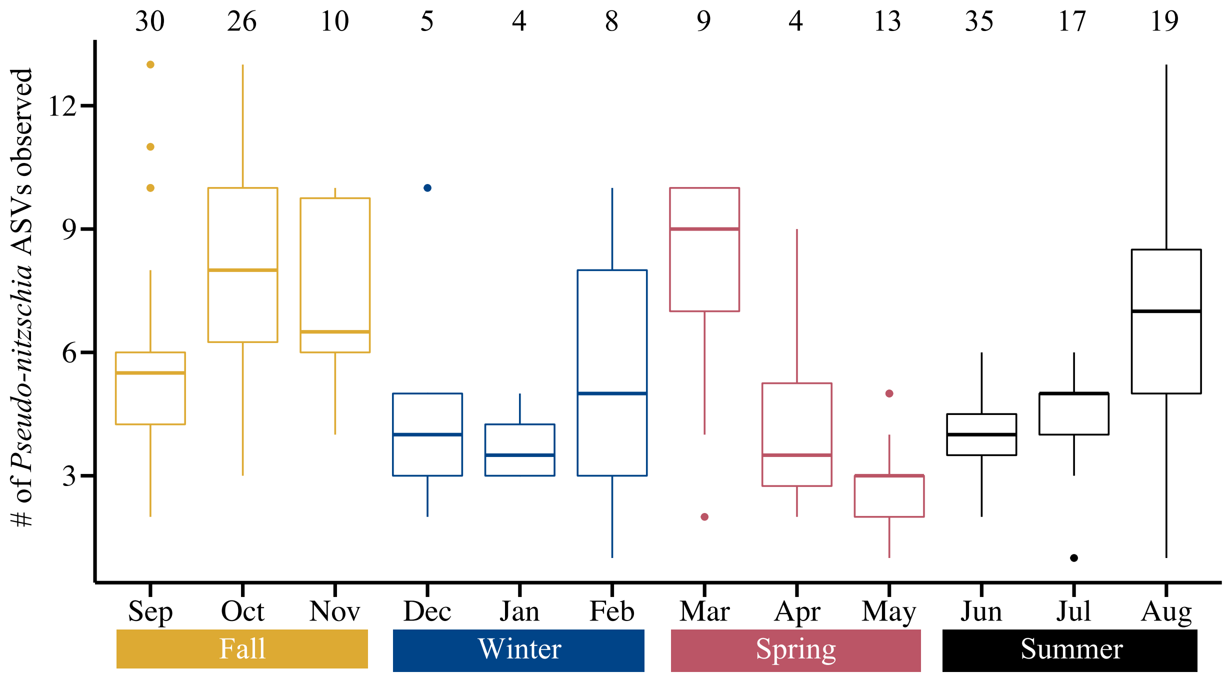


**Fig S7.** The number of *Pseudo-nitzschia* spp. amplicon sequence variants (ASVs) detected in samples (n = 180) averaged by month. This includes ASVs as present if they made-up > 1% reads by relative abundance in the particular sample. This excludes the Northeast Shelf Long-Term Ecological Research (NES LTER) cruise samples, but includes precautionary closure, closure, and net tow samples along with any samples collected by this study. The number of samples collected per month is indicated along the top by the numbers, and the seasonal groupings of the months are indicated along the bottom.

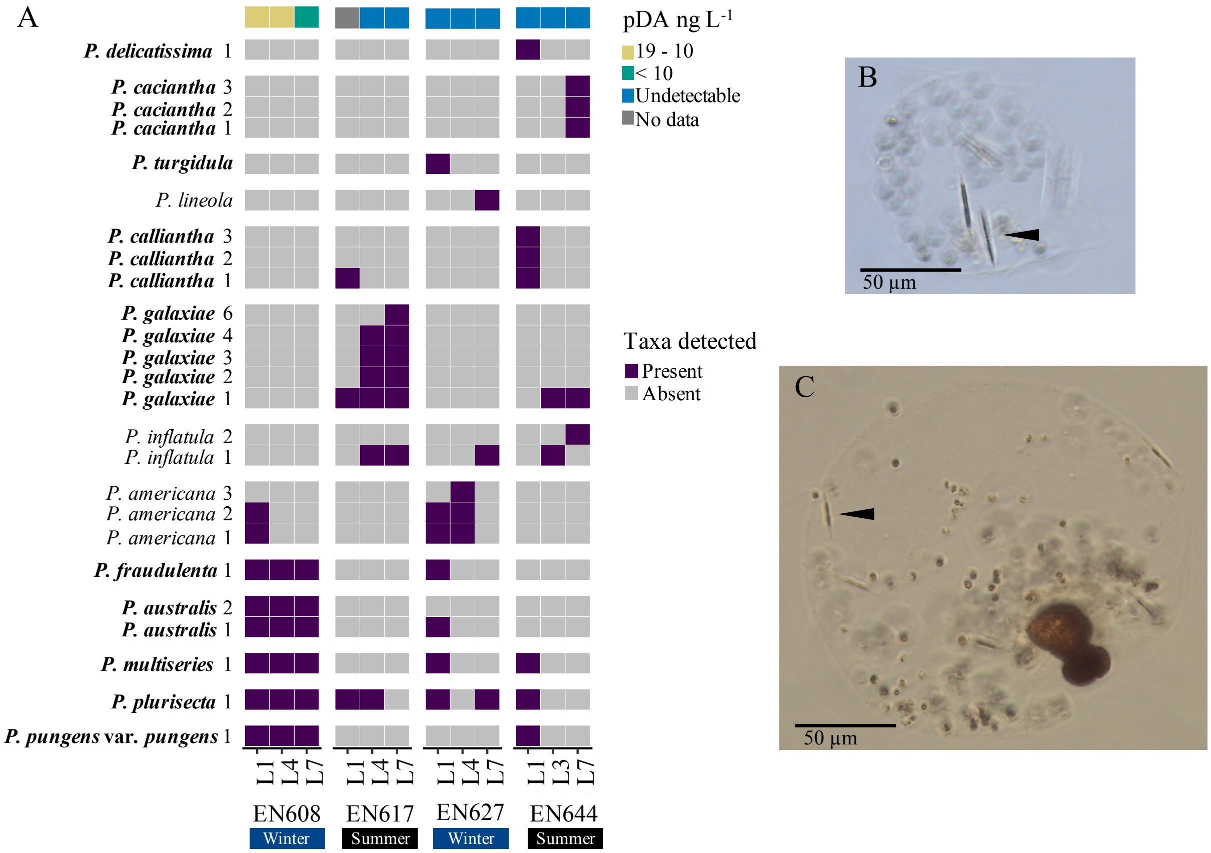


**Fig S8.** *Pseudo-nitzschia* spp. in offshore samples from the Northeast Shelf Long Term Ecological Research (NES LTER) and related stations. **[A]** Heatmap of amplicon sequence variants identified as *Pseudo-nitzschia* spp. from two-years of offshore NES LTER cruises (n = 12). Species known to produce domoic acid (DA) according to Bates *et al.* 2018 are denoted in bold. EN608 samples are from 1/31/18 – 2/4/18, EN617 was 7/20/18 – 7/22/18, EN627 was 2/1/19 – 2/3/19, and EN644 was 8/21/19 – 8/22/19. **[B]** Light microscopy image of possible *Pseudo-nitzschia* cells (arrow for example) associating with another plankton taxon from a sample taken during EN608 from the underway system near L3 at 40.9884 N, -70.8801 W (02/04/2018). This sample is fixed in Lugol's and was observed during cell counts using a Sedgewick-Rafter chamber. This sample did not have a corresponding DNA sequencing sample. **[C]** Light microscopy image of possible *Pseudo-nitzschia* cells (arrow for example) associating with other plankton taxa from a sample taken from the Martha's Vineyard Coastal Observatory (MVCO) station at 41.3223 N, -70.5760 W (03/24/2018), near the beginning of the NES LTER transect. This sample is fixed in Lugol's and was observed during cell counts using a Sedgewick-Rafter chamber. This sample did not have a corresponding DNA sequencing sample.

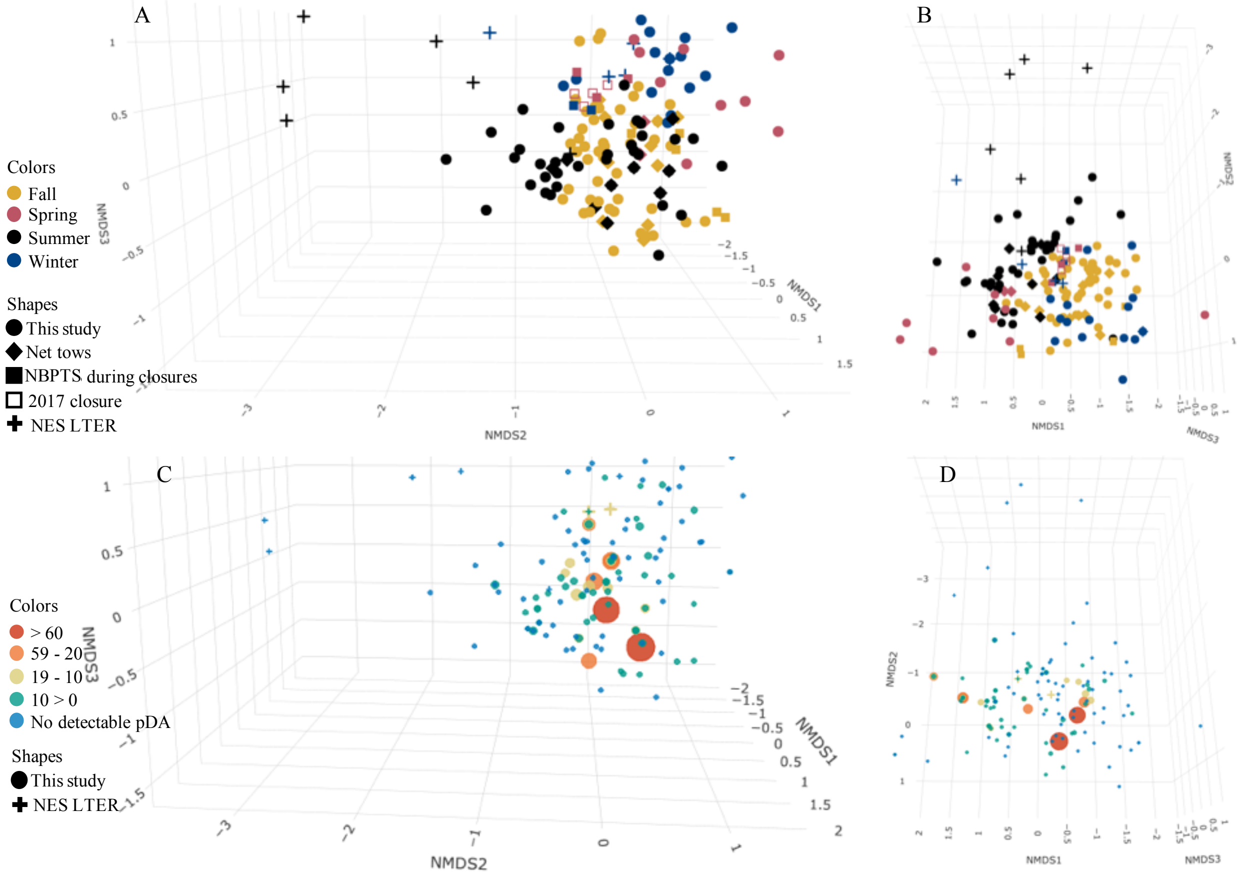


**Fig S9.** Nonmetric multidimensional scaling (NMDS) of Jaccard distance matrix of the presence or absence of *Pseudo-nitzschia* spp. amplicon sequence variants in samples from Narragansett Bay (NBay), RI and offshore research cruises (n = 192). Both plots are the same NMDS with stress = 0.0924. Solution was not reached by defaults, so iterations raised to 100 and dimensions were raised to three which these are plotted in 3D space with different views shown. **[A, B]** *Pseudo-nitzschia* spp. assemblages across seasons (color) by sample group (shape) including the Long-Term Plankton Time Series (NBPTS) and Northeast Shelf Long-Term Ecological Research (NES LTER) cruises, which have different sample types such as pore size and sample years. **[C] [D]** Assemblages of *Pseudo-nitzschia* taxa overlaid with concentration of particulate domoic acid (pDA) measured in filtered seawater $L^{-1}$. Samples without pDA measurements are not shown in the plot but included in the NMDS.

**Table S1.** Sampling sites in Narragansett Bay, Rhode Island (RI), USA and offshore.

| Site | Latitude (N) | Longitude (W) | Sampled by | Approximate frequency with date range | Access | Notes |
|---|---|---|---|---|---|---|
| Long-Term Plankton Time Series (NBPTS) | 41.57 | -71.39 | NBPTS Assistant & this study | Weekly; fall ‘17 – winter ‘19 | Boat | Most sampled in tandem with the URI GSO NBPTS; historical plankton data |
| Whale Rock (WR) | 41.34 | -71.42 | Fish Trawl Assistant & this study | Weekly; fall ’17, summer ’18 – winter ‘19 | Boat | Most sampled in tandem with the URI GSO Fish Trawl |
| East Passage (EP) | 41.45 | -71.38 | This study | Weekly; fall ’17, summer & fall ’18, spring - fall ‘19 | Boat | RI Dept. of Environmental Management (DEM) harmful algal bloom (HAB) monitoring site 14E-8 |
| Castle Hill Beach (CHB) | 41.46 | -71.36 | This study | Weekly; winter ‘18 – fall ‘19 | Shore | |
| Fort Wetherill (FW) | 41.48 | -71.36 | This study | Weekly; fall ’17, spring - fall ‘18 | Shore | RI DEM HAB monitoring site 4A-3 |
| GSO Dock (GD) | 41.49 | -71.42 | This study | Weekly; fall ‘17, spring - fall ’18, spring & summer ‘19 | Shore | Used for NBPTS if boat unable to go out |
| EN608 L1 | 41.1964 | -70.8777 | Northeast U.S. Shelf (NES) Long-Term Ecological Research (LTER) | 1x; 2018-01-31 | Boat | Cast 1, Niskin 16, 3 m |
| EN608 L4 | 40.6930 | -70.8842 | | 1x; 2018-02-01 | | Cast 8, Niskin 17, 5 m |
| EN608 L7 | 40.2060 | -70.8626 | | 1x; 2018-02-03 | | Cast 23, Niskin 1, 7 m |
| EN617 L1 | 41.20 | -70.88 | | 1x; 2018-07-20 | | Flowthrough, 5 m |
| EN617 L4 | 40.7019 | -70.8767 | | 1x; 2018-07-21 | | Cast 8, Niskin 24, 2 m |
| EN617 L7 | 40.2410 | -70.8891 | | 1x; 2018-07-22 | | Cast 13, Niskin 13, 3 m |
| EN627 L1 | 41.1981 | -70.8794 | | 1x; 2019-02-01 | | Bucket, 0 m |
| EN627 L4 | 40.6951 | -70.8768 | | 1x; 2019-02-02 | | Cast 11, Niskin 12, 3 m |
| EN627 L7 | 40.2288 | -70.8826 | | 1x; 2019-02-03 | | Cast 17, Niskin 12, 5 m |
| EN644 L1 | 41.1968 | -70.8776 | | 1x; 2019-08-21 | | Cast 1, Niskin 18, 2 m |
| EN644 L3 | 40.8632 | -70.8789 | | 1x; 2019-08-21 | | Cast 5, Niskin 16, 3 m |
| EN644 L7 | 40.2135 | -70.8844 | | 1x; 2019-08-22 | | Cast 8, Niskin 19, 4 m |
| D1 | 41.6716 | -71.3616 | RI DEM; K. Hubbard | 1x; spring ’17 on 03/13/2017 | Boat | RI DEM HAB site: 1B-3 |
| D2 | 41.6636 | -71.2416 | | | | 5A-1 |
| D3 | 41.5545 | -71.4053 | | | | 3W-16 |
| D4 | 41.5776 | -71.3073 | | | | 4A-9 |
| D5 | 41.5997 | -71.2285 | | | | 5B-1 |
| A1 | 41.5212 | -71.3982 | L. Maranda | 1x; summer ‘18 | Boat | |

| | | | |
|---|---|---|---|
| A2 | 41.4435 | -71.4110 | 3x; fall ’17 & summer ‘18 |
| A3 | 41.4333 | -71.3370 | 16x; fall – winter ’17, summer ’18, spring – fall ‘19 |
| A4 | 41.4565 | -71.3758 | 3x; summer – fall ‘18 |
| A5 | 41.5132 | -71.3480 | 1x; summer ‘18 |
| A6 | 41.5673 | -71.3228 | 1x; summer ‘18 |
| A7 | 41.5910 | -71.3845 | 1x; summer ‘18 |

**Table S2.** NCBI Accession Numbers of *Pseudo-nitzschia* spp. ITS1 sequences used in custom database for taxonomy assignment. Sequences downloaded from NCBI on April 2, 2019.

| Accession Number | Species/Strain |
|---|---|
| KC409108.1 | *Pseudo-nitzschia abrensis* NerJ2 |
| KC409109.1 | *Pseudo-nitzschia abrensis* NerJ3 |
| KR021327.1 | *Pseudo-nitzschia abrensis* Pnmi19 |
| KR021323.1 | *Pseudo-nitzschia abrensis* Pnmi81 |
| KR021325.1 | *Pseudo-nitzschia abrensis* Pnmi93 |
| EU523099.1 | *Pseudo-nitzschia americana* |
| KR053127.1 | *Pseudo-nitzschia americana* GH14 |
| DQ813840.1 | *Pseudo-nitzschia arenysensis* AL11 |
| KC409093.1 | *Pseudo-nitzschia arenysensis* NerF2* |
| KC409094.1 | *Pseudo-nitzschia arenysensis* NerJ1 |
| KR053150.1 | *Pseudo-nitzschia australis* GH28 |
| KR053152.1 | *Pseudo-nitzschia australis* GH31 |
| KR021331.1 | *Pseudo-nitzschia batesiana* Pnmi12 |
| KX572953.1 | *Pseudo-nitzschia batesiana* PnMi32 |
| KX572954.1 | *Pseudo-nitzschia batesiana* PnMi44 |
| KC147514.1 | *Pseudo-nitzschia batesiana* PnTb19 |
| KR021318.1 | *Pseudo-nitzschia bipertita* Pnmi04 |
| DQ062662.1 | *Pseudo-nitzschia brasiliana* Xt3C |
| MH376339.1 | *Pseudo-nitzschia bucculenta* L2.6 |
| DQ813834.1 | *Pseudo-nitzschia caciantha* AL56 |
| DQ990361.1 | *Pseudo-nitzschia calliantha* 45 |
| DQ990360.1 | *Pseudo-nitzschia calliantha* B3 |
| DQ530621.1 | *Pseudo-nitzschia calliantha* B4 |
| KT247437.1 | *Pseudo-nitzschia calliantha* PC06 |
| KT247439.1 | *Pseudo-nitzschia calliantha* PC101 |
| AY257855.1 | *Pseudo-nitzschia calliantha* TA1 |
| KC017463.1 | *Pseudo-nitzschia calliantha* WAG |
| DQ813827.1 | *Pseudo-nitzschia cuspidata* AL17 |
| AY257852.1 | *Pseudo-nitzschia cuspidata* Mex12 |
| AY257862.1 | *Pseudo-nitzschia cuspidata* Sydney1 |
| AY257853.1 | *Pseudo-nitzschia cuspidata* Tenerife8 |
| EU523106.1 | *Pseudo-nitzschia decipiens* |
| DQ336157.1 | *Pseudo-nitzschia decipiens* GranCan41 |
| DQ336156.1 | *Pseudo-nitzschia decipiens* Mex13 |
| AY519321.1 | *Pseudo-nitzschia delicatissima* 11301 |
| AY519322.1 | *Pseudo-nitzschia delicatissima* 11401* |
| AY519329.1 | *Pseudo-nitzschia delicatissima* 12601 |

| | |
|---|---|
| AY519330.1 | *Pseudo-nitzschia delicatissima* 13301 |
| AY519297.1 | *Pseudo-nitzschia delicatissima* 2001 |
| AY519337.1 | *Pseudo-nitzschia delicatissima* 2602 |
| AY519300.1 | *Pseudo-nitzschia delicatissima* 3001 |
| AY519301.1 | *Pseudo-nitzschia delicatissima* 3201 |
| AY519303.1 | *Pseudo-nitzschia delicatissima* 3401 |
| AY519306.1 | *Pseudo-nitzschia delicatissima* 4201 |
| AY519307.1 | *Pseudo-nitzschia delicatissima* 4301 |
| AY519312.1 | *Pseudo-nitzschia delicatissima* 5201 |
| AY519317.1 | *Pseudo-nitzschia delicatissima* 7501 |
| DQ530625.1 | *Pseudo-nitzschia delicatissima* A4 |
| DQ813829.1 | *Pseudo-nitzschia delicatissima* AL22 |
| DQ813832.1 | *Pseudo-nitzschia delicatissima* AL22 |
| DQ329210.1 | *Pseudo-nitzschia delicatissima* Castell1 |
| DQ530624.1 | *Pseudo-nitzschia delicatissima* D5 |
| KR053128.1 | *Pseudo-nitzschia delicatissima* GH16 |
| DQ329208.1 | *Pseudo-nitzschia delicatissima* OFPd972 |
| KT247428.1 | *Pseudo-nitzschia delicatissima* PD111 |
| DQ813843.1 | *Pseudo-nitzschia delicatissima* SAL1 |
| DQ336153.1 | *Pseudo-nitzschia dolorosa* 300 |
| DQ813837.1 | *Pseudo-nitzschia dolorosa* AL67 |
| DQ336151.1 | *Pseudo-nitzschia dolorosa* BP3 |
| DQ336154.1 | *Pseudo-nitzschia dolorosa* Calif3 |
| KC329502.1 | *Pseudo-nitzschia fraudulenta* Pn_1 |
| KC329503.1 | *Pseudo-nitzschia fraudulenta* Pn_8 |
| DQ990364.1 | *Pseudo-nitzschia fraudulenta* PO2 |
| JN050287.1 | *Pseudo-nitzschia fryxelliana* NWFSC242 |
| KC147520.1 | *Pseudo-nitzschia fukuyoi* PnTb55 |
| AY257850.1 | *Pseudo-nitzschia galaxiae* Mex23 |
| DQ336158.1 | *Pseudo-nitzschia galaxiae* Sydney4 |
| EU051654.1 | *Pseudo-nitzschia granii* |
| KR053148.1 | *Pseudo-nitzschia hasleana* GH11 |
| KC017468.1 | *Pseudo-nitzschia hasleana* HAWK4 |
| JN091762.1 | *Pseudo-nitzschia heimii* NWFSC205 |
| DQ329204.1 | *Pseudo-nitzschia inflatula* no7 |
| KR021307.1 | *Pseudo-nitzschia kodamae* Pnmi66 |
| KR021308.1 | *Pseudo-nitzschia kodamae* Pnmi67 |
| KR021305.1 | *Pseudo-nitzschia kodamae* Pnmi75 |
| KR021313.1 | *Pseudo-nitzschia limii* Pnmi06 |
| KR021312.1 | *Pseudo-nitzschia limii* Pnmi140 |

| | |
|---|---|
| KR021311.1 | *Pseudo-nitzschia limii* Pnmi16 |
| JN091756.1 | *Pseudo-nitzschia lineola* NWFSC188 |
| KC147523.1 | *Pseudo-nitzschia lundholmiae* PnTb10 |
| KC147524.1 | *Pseudo-nitzschia lundholmiae* PnTb21 |
| KC147526.1 | *Pseudo-nitzschia lundholmiae* PnTb28 |
| KC147529.1 | *Pseudo-nitzschia lundholmiae* PnTb48 |
| DQ813839.1 | *Pseudo-nitzschia mannii* AL101 |
| DQ813842.1 | *Pseudo-nitzschia mannii* CAL1 |
| KC017449.1 | *Pseudo-nitzschia micropora* PS90/2 |
| DQ062664.1 | *Pseudo-nitzschia multiseries* OFPm984 |
| EU302796.1 | *Pseudo-nitzschia multiseries* PM02 |
| DQ990367.1 | *Pseudo-nitzschia multistriata* CM1 |
| DQ990368.1 | *Pseudo-nitzschia multistriata* CM2 |
| DQ990369.1 | *Pseudo-nitzschia multistriata* CM3 |
| KT247442.1 | *Pseudo-nitzschia multistriata* PSM11 |
| KT247443.1 | *Pseudo-nitzschia multistriata* PSM11 |
| KT247444.1 | *Pseudo-nitzschia multistriata* PSM11 |
| KT247441.1 | *Pseudo-nitzschia multistriata* PSM11 |
| MG787879.1 | *Pseudo-nitzschia nanaoensis* MC4206 |
| MH376351.1 | *Pseudo-nitzschia plurisecta* L1.4 |
| MH376350.1 | *Pseudo-nitzschia plurisecta* L3.9 |
| FJ859051.1 | *Pseudo-nitzschia pseudodelicatissima* 10A3 |
| FJ859047.1 | *Pseudo-nitzschia pseudodelicatissima* 8A9 |
| DQ813826.1 | *Pseudo-nitzschia pseudodelicatissima* AL15 |
| DQ813831.1 | *Pseudo-nitzschia pseudodelicatissima* AL29 |
| DQ813833.1 | *Pseudo-nitzschia pseudodelicatissima* AL41 |
| DQ813836.1 | *Pseudo-nitzschia pseudodelicatissima* AL60 |
| AY544769.1 | *Pseudo-nitzschia pungens* |
| DQ166533.1 | *Pseudo-nitzschia pungens* |
| KR053156.1 | *Pseudo-nitzschia pungens* PC39 |
| KR053157.1 | *Pseudo-nitzschia pungens* PC43 |
| KC329504.1 | *Pseudo-nitzschia pungens* Pn_C5A |
| KT247433.1 | *Pseudo-nitzschia pungens* PP071 |
| EU684236.1 | *Pseudo-nitzschia pungens* var. *aveirensis* Theta4 |
| MH376355.1 | *Pseudo-nitzschia pungens* var. *cingulata* L3.13 |
| KC409100.1 | *Pseudo-nitzschia pungens* var. *pungens* NerJ9 |
| KC409101.1 | *Pseudo-nitzschia pungens* var. *pungens* NerL1 |
| KM400611.1 | *Pseudo-nitzschia sabit* PnPd75 |
| KM400609.1 | *Pseudo-nitzschia sabit* PnPd76 |
| KP288507.1 | *Pseudo-nitzschia sabit* Ps147 |

| | |
|---|---|
| KP288508.1 | *Pseudo-nitzschia sabit* Ps149 |
| KP288505.1 | *Pseudo-nitzschia sabit* Ps277 |
| DQ062667.1 | *Pseudo-nitzschia seriata* f. *obtusa* T5 |
| DQ062666.1 | *Pseudo-nitzschia seriata* Lynaes8 |
| MF374771.1 | *Pseudo-nitzschia simulans* MC940 |
| MF374772.1 | *Pseudo-nitzschia simulans* MC984 |
| DQ329205.1 | *Pseudo-nitzschia subcurvata* 1F |
| KR021295.1 | *Pseudo-nitzschia subfraudulenta* Pnmi188 |
| KR021299.1 | *Pseudo-nitzschia subfraudulenta* Pnmi71 |
| KR021301.1 | *Pseudo-nitzschia subfraudulenta* Pnmi82 |
| EU523104.1 | *Pseudo-nitzschia subpacifica* |
| JN091764.1 | *Pseudo-nitzschia turgidula* NWFSC220 |
| AY257839.1 | *Pseudo-nitzschia turgiduloides* 319 |

*Indicates identical sequences
